# Thresholding Gini Variable Importance with a single trained Random Forest: An Empirical Bayes Approach

**DOI:** 10.1101/2022.04.06.487300

**Authors:** Robert Dunne, Roc Reguant, Priya Ramarao-Milne, Piotr Szul, Letitia Sng, Mischa Lundberg, Natalie A. Twine, Denis C. Bauer

## Abstract

**Background:** Random Forests (RF) are a widely used modelling tool, enabling feature-selection via a variable importance measure. For this, a threshold is required that separates label-associated features from false positives. In the absence of a good understanding of the characteristics of the variable importance measures, current approaches attempt to select features by training multiple RFs to generate statistical power via a permutation null, employ recursive feature elimination or a combination of both. However, for high-dimensional datasets, such as genome data with millions of variables, this is computationally infeasible.

**Method:** We present RFlocalfdr, a statistical approach for thresholding that identifies which features are significantly associated with the prediction label and reduces false positives. It builds on the empirical Bayes argument of Efron (2005) and models the variable importance as mixture of two distributions – null and non-null “genes.”

**Result:** We demonstrate on synthetic data that RFlocalfdr has an equivalent accuracy to computationally more intensive approaches, while being up to 100 times faster. RFlocalfdr is the only tested method able to successfully threshold a dataset with 6 Million features and 10,000 samples. RFlocalfdr performs analysis in real-time and is compatible with any RF implementation that returns variable importance and counts, such as ranger or VariantSpark.

**Conclusion:** RFlocalfdr allows for robust feature selection by placing a confidence value on the predicted importance score. It does so without repeated fitting of the RF or the use of additional shadow variables and is thus usable for data sets with very large numbers of variables.

## 1 Introduction

Random Forests (RF) is a non-linear modelling tool that has widespread popularity, from health care to academia to industry (Lundberg et al., 2019). This prevalence is, in part, due to the ability of RF to process large volumes of data efficiently (Bayat et al., 2020), which is especially useful for genetic data which are high-dimensional in nature with *p* >> *n*. Other RF features beneficial for genetic data include robust default hyperparameters and the use of out-of-bag (OOB) estimates of true error in place of a test/train data split, which can be difficult with small sample sizes (Janitza and Hornung, 2018). Crucially, RF provide variable-importance-measures (VIM) which enables the selection of features associated with the prediction label via a threshold value and in most genetic data applications, is the desired outcome. As such, RF are a step in the direction of interpretable machine learning.

However, VIMs are vulnerable to false positives and a statistical assessment is needed to confidently identify which variables are significantly label associated. Although there is no theoretically defined VIM in the sense of a parametric quantity that a variable importance estimator should try to estimate (Grömping 2009), there are two with strong theoretical foundations: Shapley values (Lundberg and Lee, 2017; Lundberg et al., 2019, 2020)) and “conditional variable importance” (Strobl et al., 2008). However, for wide data like genomic data, the computational burden to generate these VIM is too high for practical use.

Shapley values have a strong theoretical foundation (Lundberg and Lee, 2017; Lundberg et al., 2019, 2020)) and “conditional variable importance” (Strobl et al., 2008) overcomes some of the bias problems of RF VIM. However, for wide data like genomic data, the computational burden to generate these VIM is too high for practical use.

Nevertheless, there are two broad approaches to determining the label-association for RF VIM, permutation of the response vector and recursive-feature-elimination (RFE) (Degenhardt et al., 2019). With the permutation approach, the response vector is permuted *k* times to calculate *p* values for significance from the permuted VIM, akin to a standard procedure for estimating false discovery rates. However, the validity of this procedure relies on the assumption that the statistic for gene *j* is a function of only the data for gene *j*, and as the VIMs are functionally related, this assumption is violated, making a permutation approach inappropriate for genomic data (Witten and Tibshirani, 2008; Huynh-Thu et al., 2012). The actual-impurity-reduction (AIR) (Nembrini et al., 2018) in combination with the Vita approach (Janitza et al., 2016), and the Permutation IMPortance (PIMP) (Altmann et al., 2010) algorithms rely on such a permutation approach to calculate *p*-values and in the case of PIMP, multiple testing correction should be applied. The second widely used method is RFE where RF are built recursively, removing a proportion of least important features before a new RF is generated with the remaining variables until a single feature is left. The verSelRF algorithm (Díaz-Uriarte and Alvarez de Andrés, 2006) and its modifications (Degenhardt et al., 2019), uses the RFE approach. Combining permutation and RFE approaches, Boruta (Kursa and Rudnicki, 2010) generates shadow features by permuting the original features and recursively trains RF on this extended set until a stopping criterion is met. A test comparing the VIM of the real features are compared to the maximum of all the shadow features to determine which features are removed.

As some of these feature selection approaches (i.e., RFE, PIMP, AIR) require multiple RF to be trained, they are computationally intensive and prohibitive for high-dimensional data like genetic datasets which scale up to 500,000 samples × > 90 million genotypes (Bycroft et al., 2018). Even with a smaller dataset of 500 samples ×50000 gene expression levels (for example, the colon cancer data set of LaPointe et al. (2012)) with, say, *k* = 100 genes that are highly involved with the label, we can define *ρ* = *k*/*n* and *δ* = *n*/*p* to get {*ρ, δ*} = {0.2, 0.01}, putting it in a “difficult” region of {*ρ, δ*} phase space where many algorithmic methods of feature recovery have a high probability of failing Donoho and Stodden (2006); Donoho and Tanner (2009)

From a statistical standpoint, *p* simultaneous tests can be performed on a genetic dataset 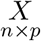 with *p* >> *n* to get the test statistics 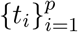. Under assumptions about the distribution 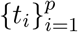, statistics *z*_*j*_ can be computed, comparing cases with controls which should give, by the central limit theorem, *z*_*j*_ ∼ *N* (Δ_*j*_, 1), where Δ_*j*_ is the effect size for gene *j*. Therefore, |Δ_*j*_| is small for “null genes” (i.e., genes that show the same activity in cases and controls), while |Δ_*j*_| is large for genes having much different responses for cases versus controls. Inference for the individual *p* genes gives the *p*-value, 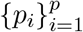.

This is the position taken by the widely used LIMMA (linear models for microarray data) approach (Ritchie et al., 2015). By using an empirical Bayes argument to make an adjustment to the variance of each gene based on a model for the variance of all genes in the sample, genes with very small variance are prevented from having a greatly inflated *t* statistic and from appearing significant. Another common practice is to permute the phenotype and derive a matrix of statistics of dimension *p* genes by *M* permutations. Approaches to determining the threshold of significance given such a matrix are discussed in Churchill and Doerge (1994).

For both approaches, the issue of multiple testing needs to be evaluated and there are three widely used approaches in genetic studies: false discovery rate (FDR) Benjamini and Hochberg (1995), Bonferroni correction (Korthauer et al., 2019) 2019), and q-value (Storey and Tibshirani, 2003). The FDR approach has demonstrated greater power to detect true positives than the simpler Bonferroni correction, while the *q*-value builds on the infimum over the *p*-values of the FDR. Further to this, the “local FDR” approach was proposed to control the FDR using an empirical Bayes estimate of the (Efron (2005; 2007; 2008; 2010).

To address the shortcomings of currently available feature selection approaches including AIR and Boruta, we introduce RFlocalfdr, a method for setting a significance level of the VIM, mean decrease in impurity (MDI) importances, based on Efron’s local FDR approach. The RFlocalfdr approach does not involve any refitting of RF and does not use “shadow variables” (Nembrini et al., 2018), making it applicable to extremely high dimensional datasets, including genomics where the number of features may be in the millions.

## 2 Methods

### 2.1 Illustration of Efron’s Empirical Bayes Approach

There are several points that make an ideal situation for an empirical Bayes estimate of the null distribution as discussed in Efron (2010). Firstly, the dataset is composed of two groups: a large group of data that will generate null values of some statistic *z*, and a much smaller group that will generate non-null values. This means that we have sufficient data to model the null distribution, which Efron argues we should always do in cases like this, rather than make a distributional assumption. By adopting this approach, the high dimensionality of genetic datasets is now an asset as there are enough data points to estimate the null distribution accurately. Secondly, despite the large “sample” sizes, the *N* (0, 1) Gaussian distribution be seen to have a very poor fit to the *z* values.

We illustrate Efron’s empirical Bayes approach using a dataset from Hedenfalk et al. (2001), consisting of a matrix with 3,226 rows corresponding to the expression levels of genes and 15 columns corresponding to the 15 samples, divided between tumours with the BRCA1 and BRCA2 mutations. Let *t*_*i*_ be the standard *t*-statistics arising from the comparisons of cases and controls (i.e., tumours with or without the BRCA1/BRCA2 mutations). Let *z*_*i*_ = Φ^−1^(*G*_0_(*t*_*i*_)), where Φ is the standard normal cdf, and *G*_0_ is a putative null cdf for the *t*-values. *G*_0_ can be a theoretical null distribution, or a permutation null. Interestingly, in this case, the permutation density is very similar to the theoretical *N* (0, 1) density so a non-parametric approach does not alleviate the problem.

As per Efron’s approach, a histogram of the 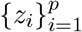 is plotted (Figure 1b) and it demonstrates that the modelling assumptions are inaccurate as the distribution of *z*_*i*_ is not a *N* (0, 1), as shown in red. Efron discusses some reasons behind this occurrence including failed assumptions, correlations between cases or between features, and unobserved covariates.

**Figure 1:**
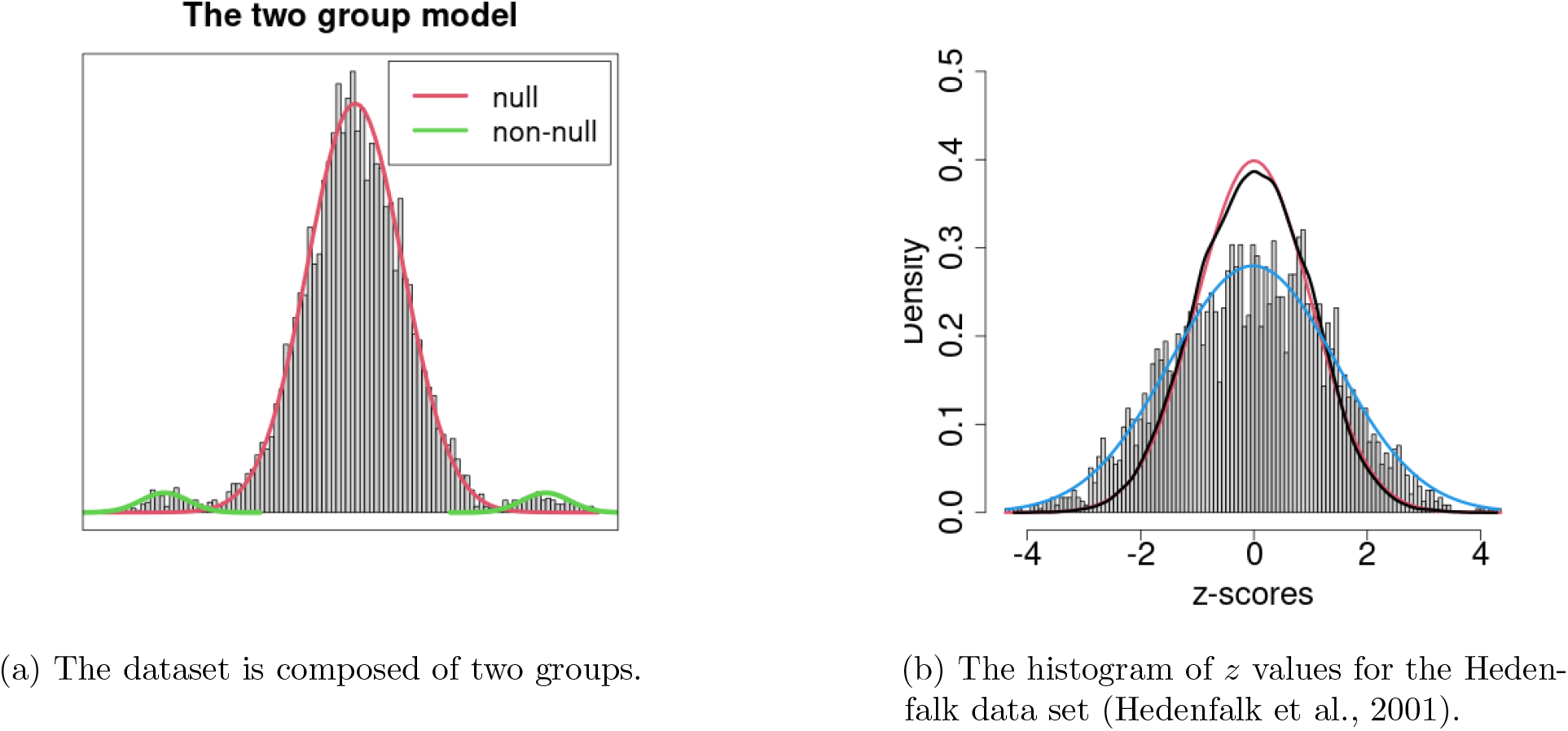
(a) The two group model. We model the data as generated by two processes, one of which produces a set of null statistics (density shown in red) and one which produces non-null statistics (shown in green). (b) The histogram of *z* values for the Hedenfalk data set (Hedenfalk et al., 2001). The red curve shows the *N* (0, 1) distribution and the black curve shows the permutation null distribution, which is similar to the theoretical *N* (0, 1) curve. The blue curve is the empirical Bayes Gaussian fit to the data.

The observed distribution is thus modelled as a mixture *f* (*z*) = *p*_*o*_*f*_*o*_(*z*) + (1 − *p*_0_)*f*_1_(*z*) where *f*_*o*_(*z*) is the null distribution of *t*-statistics and *f*_1_(*z*) is the distribution of significant *t*-statistics (Figure 1). The modelling process involves using the central mass of the null distribution to fit a Gaussian and then the local FDR is calculated as the ratio of a null density and the observed density of the tails *f*_1_(*z*). See Efron (2008, 2005, 2007, 2008, 2010)) for more details.

### 2.2 Empirical Bayes for MDI Importances

The RFlocalfdr, inspired by Efron’s approach as introduced above, is a method for modelling the distribution of Mean Decrease in Impurity (MDI) importances from RF with a view to setting a significance threshold. This method depends on having a large number of features and a large number of trees making it ideally suited for genetic analyses, including differential gene expression and GWAS.

We illustrate the RFlocalfdr method using a vector of MDI importances calculated from a RF of the 1000 Genomes Projects (Fairley et al., 2020), described further in the Results section. The density of the log transformed vector of MDI importances is considered (Figure 2a) and, as is often the case, it is multi-model. We can model this as a mixture,

**Figure 2:**
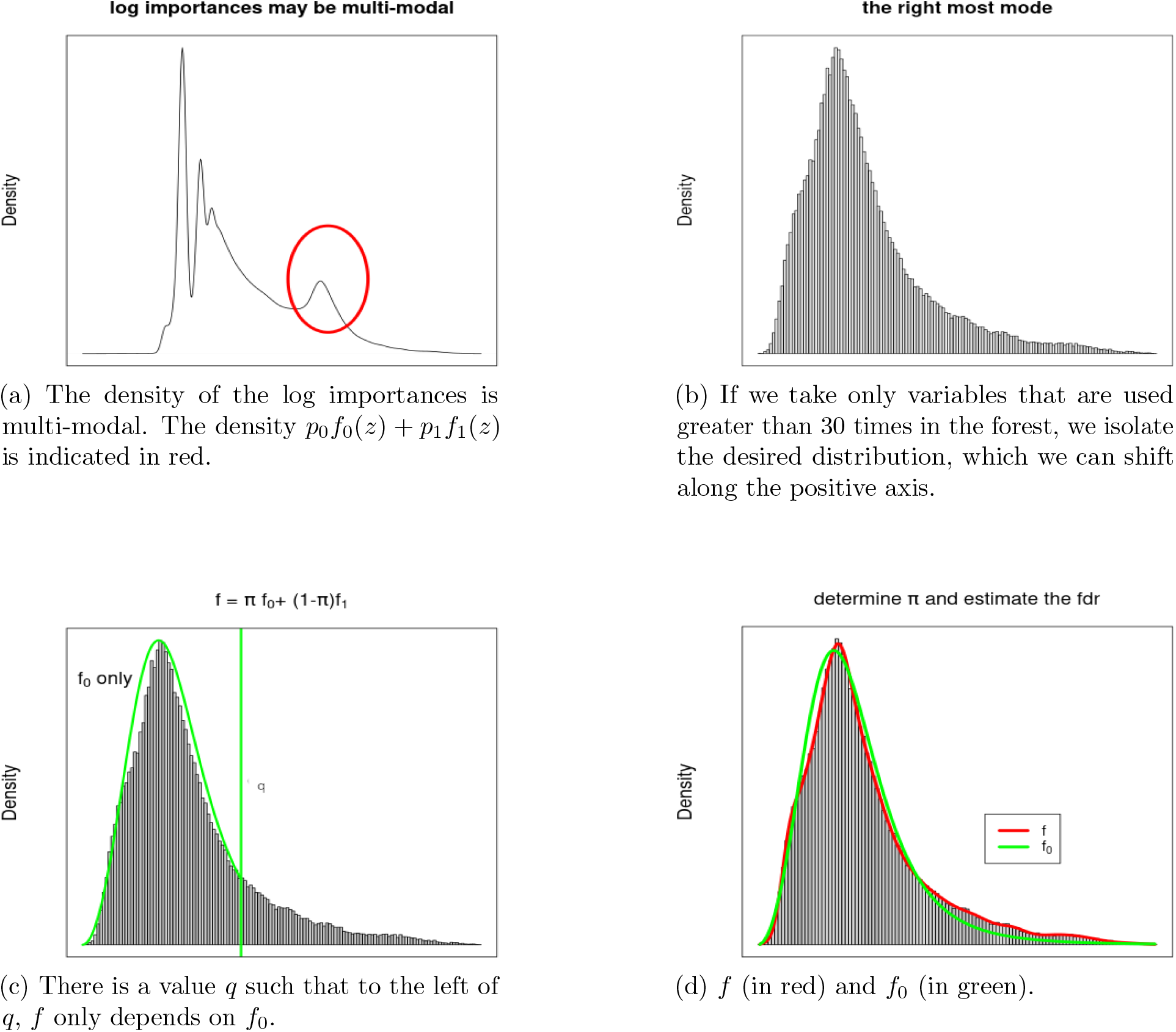
The steps in estimating the local fdr for RF MDI importance values.

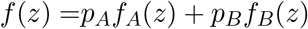

And

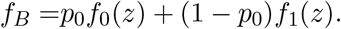

The distribution we are interested in, *f*_*B*_(*z*) = *p*_0_*f*_0_(*z*) + (1 − *p*_0_)*f*_1_(*z*), indicated in red in Figure 2a, is assumed to be unimodal and a mixture of null features and non-null features. Resultingly, we have two tasks:

- is to separate *f*_*A*_(*z*) and *f*_*B*_(*z*),
- like Efron’s problem, to estimate the empirical null *f*_0_(*z*) and calculate the local FDR.

For task 1, modelling *f*_*A*_(*z*) + *f*_*B*_(*z*) by mixtures is difficult due to the wide range of distributions between different data sets. However, the modes of *f*_*A*_(*z*) are observed to be associated with features that were used a specific (and small number) of times in the RF. For example, the left most peak of *f*_*A*_(*z*) in Figure 2a is composed of features that were used only once in the RF. There will often be another peak (at −*∞* in Figure 2a as we have taken the log) of features that were not used at any time in the RF. Building on these observations, we denote the number of times each variable is used as *C*. We then progressively threshold *C* giving *f*_*c*_(*z*) = *f* (*z*)|*C* > *c*. A skew-normal distribution (Azzalini, 2022) was explored to be a fit for *f*_*c*_(*z*), and either a Hartigan’s diptest for unimodality or a goodness-of-fit test such as the Cramer-von Mises test is applied. However, as none of these procedures selects a *c* value giving a satisfactory fit (See Web Info), our current procedure is to fit a skew-normal *S*_*q*_(*z*) up to the *q*^*th*^ quantile of *f*_*c*_(*z*), and calculate the *L*_*∞*_ norm *d*_*c*_ = max_*z*_ |*f*_*c*_(*z*) − *S*_*q*_(*z*)|. The *d*_*c*_ is plotted against *c*, and the minimum value *c*^*^ is chosen. The corresponding distribution, *f*_*c*_* (*z*), is shifted along the *z* positive axis so that the smallest value is 0. For this dataset, the selected distribution is shown in Figure 2b.

From a histogram of the selected distribution resulting from task 1 (Figure 2b), we start task 2 by fitting a spline to the observed bin counts, denoted as *f* and shown in green in Figure 2d. This can be done using standard Poisson generalised linear modelling software, fitting the counts to a natural cubic spline basis on the midpoints of the bins. By assumption, there is a point *q* such that to the left of *q f*_*B*_ ∼ *f*_0_(*z*), that is, there is a *q* such that there are only null features to the left of *q*. A change point method related to penalised model selection(Gauran et al., 2018) is used to determine *q* (Figure 2c) and then *p*_0_ is estimated. Using only the data < *q*, a skew-normal is fit to *f*_0_ using only the truncated range with the formulation of the skew-normal given by Ashour and Abdel-hameed (2010). The fit is done with non-linear least squares (Elzhov et al., 2022). With *f, p*_0_, and *f*_0_, the local fdr is estimated as 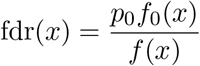 and shown in figure 3.

**Figure 3:**
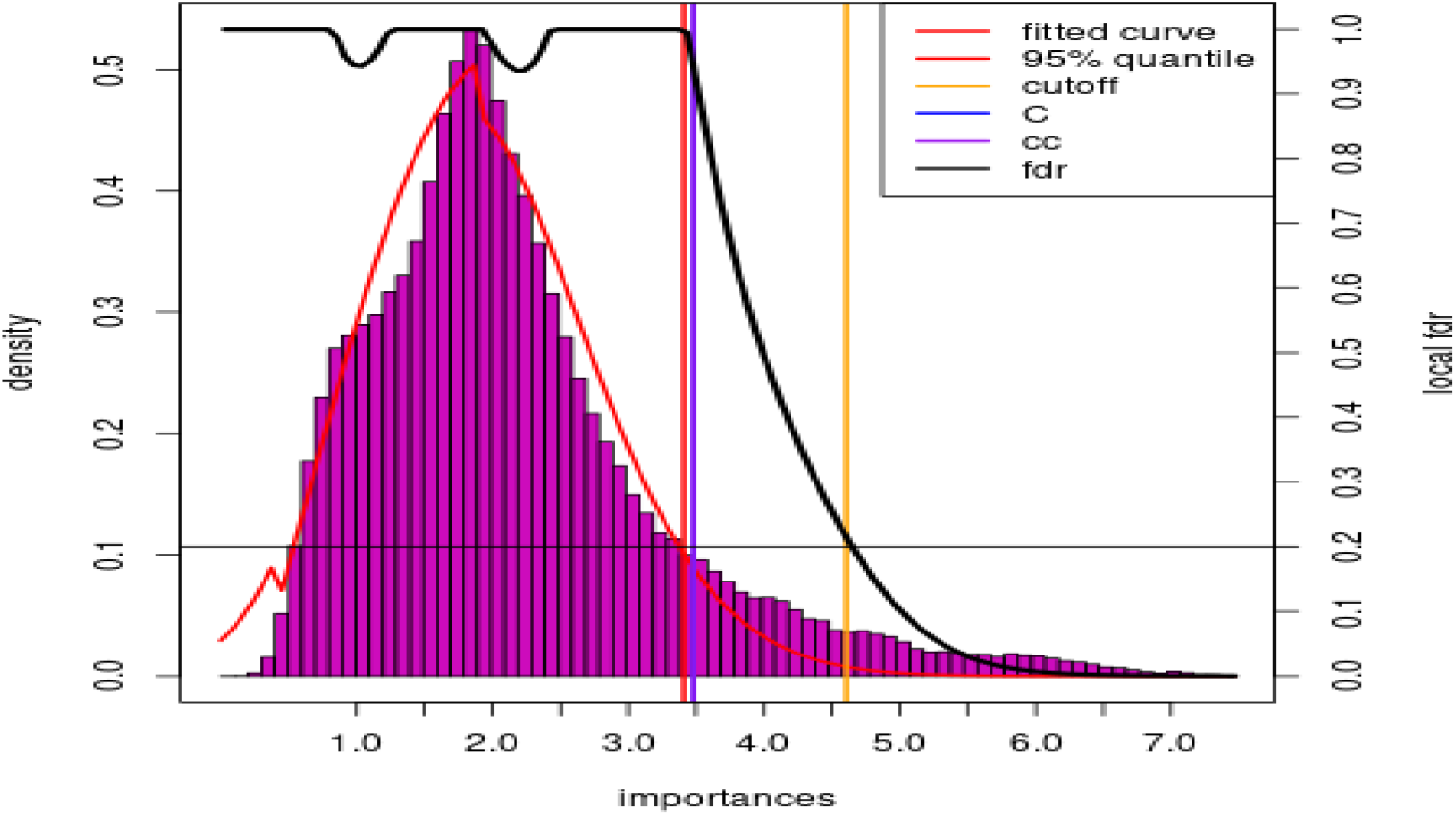
A plot produced by the code RFlocalfdr. The fdr curve is shown in black.

## 3 Results

We applied the RFlocalfdr approach on three datasets:

1. a synthetic dataset with a strong correlation structure to establish the need for statistical approaches in feature selection,
2. chromosome 22 from the 1000 Genomes Project dataset to demonstrate real-world suitability,
3. a dataset of 10^6^ data-points to show applicability to large-scale genomic datasets like whole genome sequencing.

Where possible, the RFlocalfdr approach was compared to four published approaches introduced in the previous section: AIR, Boruta, RFE, and PIMP. The R package ranger (Wright and Ziegler, 2017) was used to fir all models and the parameters used are given in the supplementary files included in the Web Information.

### 3.1 Feature Selection on a Synthetic Dataset

The synthetic dataset consists of “bands” with “blocks” of of {1, 2, 4, 8, 16, 32, 64} of identical features (Figure 4). The features are *∈* {0, 1, 2}, a common encoding for genomic data where the numbers represent the number of copies of the minor allele. Only band 1 is used to calculate the *y* vector, and *y* is 1 if any of *X*[, *c*(1, 2, 4, 8, 16, 32, 64)] is non-zero. The result of this is that *y* is unbalanced, containing more 1’s than 0’s. In total, there are 50 bands and 200 observations, so *X* is 300 × 6350 with 127 non-null features (See Web Info for more details). A standard RF was fitted to this dataset and the resulting MDI importances was recorded.

**Figure 4:**
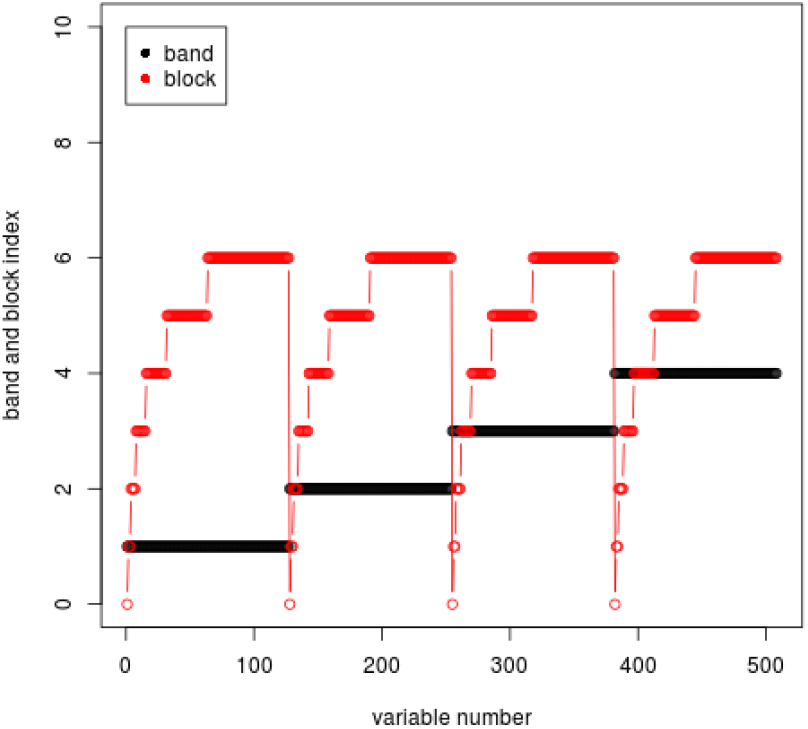
The data is structured into bands and blocks. The color and the *y*-axis show which band/block each variable belongs to, not the variables value. Each “band” contains “blocks” of sizes 1, 2, 4, 8, 16, 32, and 64. Each block consists of correlated (identical variables), where each variable is *∈* {0, 1, 2}. The dependent variable *y* is 1 if any of *X*[, *c*(1, 2, 4, 8, 16, 32, 64)] is non-zero, so only band 1 has a relationship to the dependent variable.

As shown in Figure 5, selecting features above a single importance threshold did not achieve a perfect separation of the true positive features from band one (in red) and the false positives from the other bands. This issue was further exacerbated by blocks with fewer features having higher MDI importances as the importances was “smeared” over the correlated (in this case, identical) features. The effects of correlation on MDI importances are further discussed in Web Info. Therefore, a statistical approach is necessary for true positive selection.

**Figure 5:**
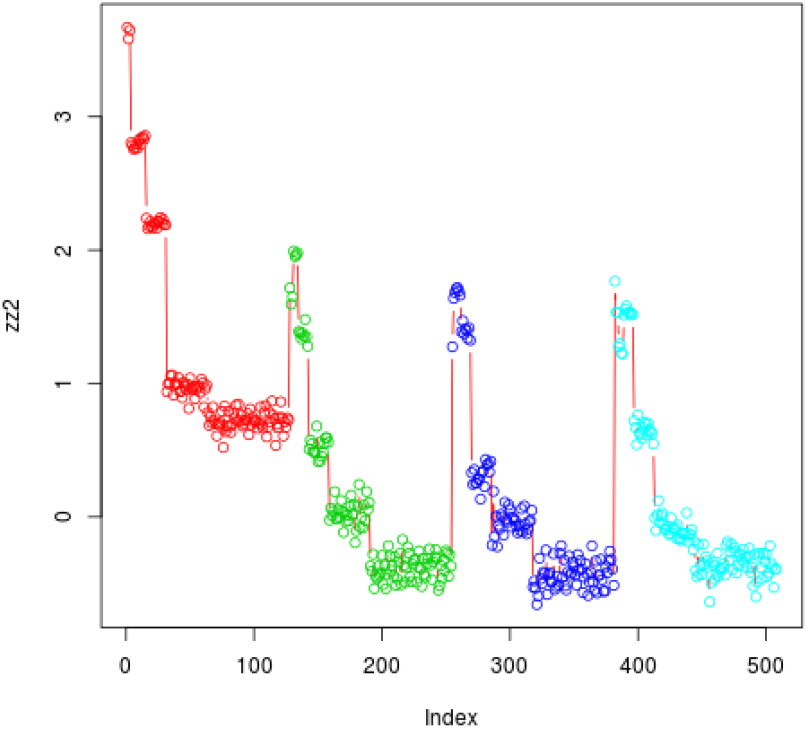
The log importance scores for the simulation, arranged by variable number and colored by band. Only the red variables are non-null, so we can see that it is not possible to threshold the importance scores so as to only recover the non-null variables.

Table 1 shows the performance measures of applying the four feature selection methods and our approach to this dataset for feature selection. Our RFlocalfdr approach resulted in the second highest AUC after AIR (0.73 vs 0.98 respectively), but unlike AIR, it also has a high sensitivity (0.99 vs 0.96 respectively). Furthermore, while Boruta and RFlocalfdr resulted in the lowest AUCs, both performed comparably on sensitivity and had the lowest number of true positives (2 and 22 respectively). See Web Info for further details, including a multiple testing correction for the PIMP values.

**Table 1:** Performance of variable selection for the simulated example by several methods. The best outcomes for each category are in bold face.

| Method | true positives | false positives | Sensitivity | AUC |
| --- | --- | --- | --- | --- |
| RFlocalfdr | 59 | 36 | **0.99** | 0.73 |
| Boruta | 2 | 2 | **0.99** | 0.50 |
| RFE | 22 | 2 | **0.99** | 0.58 |
| AIR | 127 | 273 | 0.96 | **0.98** |
| PIMP | 39 | 556 | 0.91 | 0.61 |

### 3.2 Feature Selection on the 1000 Genomes Project Dataset

The 1000 Genomes Project dataset was obtained as VCF files from their FTP site with each VCF file containing the genotypes of single nucleotide polymorphisms (SNPs) for every individual. There are 2,504 individuals with available genotypes in total and no additional processing was performed. An RF was used to predict the ethnicity of each individual using 1 million SNPs from chromosome 22. This dataset was used to illustrate the method in the previous section and the script detailing the analysis in depth can be found in Web Info.

The RFlocalfdr approach selected 6,335 SNPs that were significantly associated with ethnicity at an FDR of 0.2. Similarly, the Boruta algorithm returned 6,773 significant SNPs with a 82% overlap to RFlocalfdr (Figure 2). Unlike RFlocalfdr which offers a continuous *p*-value scale for the adjustment of FDR, Boruta offers two levels and at the most conservative “confirmed” option, it returns 1,443 SNPs. All “confirmed” SNPs were included also significant with RFlocalfdr and their RFlocalfdr p-values clustered at the lower end, suggesting higher association. Furthermore, a one-way ANOVA showed a significant association (*p* < 2 × 10^−16^) between RFlocalfdr *p*-values and Boruta categories.

As in the simulated example, AIR selects an order of magnitude (*≈* 10×) more significant SNPs than RFlocalfdr and Boruta (Table 2), potentially having a higher sensitivity. However, with the *p*-value of all 61,092 SNPs receiving a score of 0, AIR does not offer any option for tuning the specificity and the resulting false positive rate may be substantial (Figure 6). The final feature selection approach RFE appeared to have focused on specificity and returned 59 nested sets, with a set of 12 SNPs having the smallest prediction error.

**Table 2:** The number of SNPs returned by each methods and the run time. Run times are expressed in multiples of the run time of a single ranger (Wright and Ziegler, 2017) fit for the given hardware configuration. The run-time hence describes the number of “refits” that each method requires. The processing time outside of the RF fit is negligible, by comparison, in all cases.

| Method | SNPs returned | run time |
| --- | --- | --- |
| RFlocalfdr | 6,335 | 1 |
| Boruta | 6,773 | 100 |
| RFE | 12 | 50-60 |
| AIR | 61,092 | 1.5-2 |
| PIMP | 787502 | 100 |

**Figure 6:**
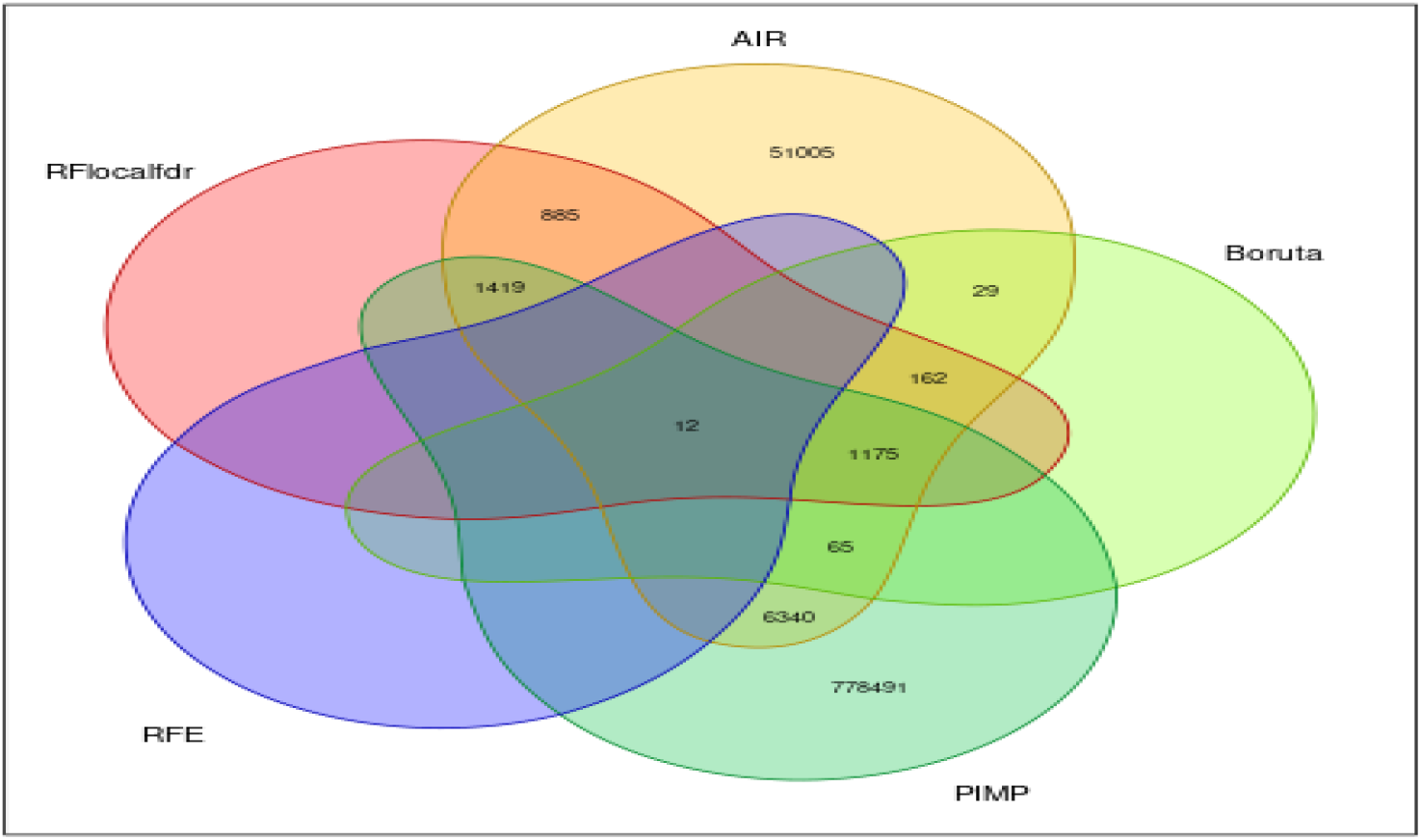
A Venn diagram of the overlaps in SNP for RFlocalfdr, Boruta, AIR, RFE and PIMP. AIR and PIMP are very much the outliers, with more than 10^4^ unique SNPs (not found by the other methods) for each method.

The run-times of each approach were evaluated using the same processing parameters next (Table 2). As RFlocalfdr does not require multiple passes of RF to be conducted, the run-time is essentially the run-time for the RF model. Further, this can be run in parallel so the run-time of the RFlocalfdr approach also depends on the number of cores being used. Expectedly, RFlocalfdr was 25 times faster than the next best performing method AIR, 57 times faster than RFE, and 100 times faster than Boruta and PIMP.

Run-time is discussed in the Web Info, where full details are given. Here we simply note:

- the number of evaluations of RF is a user set parameter for Boruta and PIMP;
- for RFE, it is data dependant and each run of RF will have a reduced number of variables and so will be faster;
- AIR requires only one run of RF but considers twice the number of variables (due to the “shadow variables”) which produces a longer run-time.

### 3.3 Feature Selection on Large-Scale Dataset (10^6^ Datapoints)

This dataset was presented in Bayat et al. (2020) with 10,000 samples and 6 million SNPs as features. Although based on the 1000 Genomes Project dataset, it has a synthetic *y* value. Due to ranger’s inability to fit RF with the size of the dataset, Apache Spark-based RF implementations VariantSpark (Bayat et al., 2020) and ReForest (Lulli et al., 2017) were used instead. ReForest fitted the model in about 15 hours, compared to VariantSpark which took 5 hours.

Using MDI and the counts reported by VariantSpark, RFlocalfd selected 38 SNPs as significant, including the 5 SNPs used to generate the simulated *y* value This resulted in a 100% success rate in recovering the significant variables with a 87% false discovery rate. Given the run-times reported in Table 2, fitting this large-scale dataset would require between 10 hours to 62 days for the other feature selection algorithms to complete. While a 10 hour estimated run time for AIR would be acceptable, neither VariantSpark nor ReForest offer this option. Hence, RFlocalfdr is the only approach capable of selecting variants associated with a trait using whole-genome size datasets in a feasible time-frame.

## 4 Discussion

In this paper, we present RFlocalfdr, a method for calculating the significance threshold of RF MDI importances for detecting label-associated features using an empirical Bayes approach. The accuracy of RFlocalfdr was shown to be comparable to other published techniques in terms of performance metrics but demonstrated advantages as well. This includes:

1. computational efficiency as it requires only a single fit of RF particularly in comparison to RFE permutation methods such as PIMP,
2. broad applicability to any RF implementation that returns MDI importances and counts of variables use, and
3. it provides a range of *p*-values allowing for tailored sensitivity and specificity selections.

Of the other methods tested in this paper, only RFE offers a similar capability to adjust the sensitivity/specificity trade-off through the selection of other nested sets with a larger or smaller number of associated features. In contrast, Boruta and AIR do not offer this capability as there is no criteria for sub-setting the set of associated features, and in practice, AIR tends to report a vastly larger number of significant features. Whether this large number of reportedly significantly features is desirable would depend on the context of the dataset, subsequent analysis, and the false discovery tolerance.

Although we have successfully processed the large-scale dataset of 106 datapoints using ranger, several hundred cores and several terabytes of RAM was required. Therefore, features selection approaches that require multiple refits along with RF implementations where run-times exponentially increase with data size will eventually become infeasible. This is particularly true for genetic analyses as availability of large-scale genomic datasets become commonplace. The RFlocalfdr approach circumvents these challenges as it only requires a single RF fit and can be applied to any RF implementation that returns MDI importances and counts of features used, such as the highly scalable VariantSpark implementation.

We note that Efron’s local FDR method is justified on its clearly superior modelling of the histogram of statistics, and not on its computational speed. Further, we have made no claim about the utility of RFlocalfdr, merely sown that it is comparable to other feature selection methods and is computationally much faster. However, while the MDI importances offer a number of challenges that are not present in the case/control comparison that Efron’s local FDR approach was designed for, it is conceivable that it may have better properties than other methods of feature selection. More experience in the application of RFlocalfdr would be required to ascertain this.

Early applications of this method have led to promising results where the use of RFlocalfdr in tandem with VariantSpark and BitEpi (Bayat et al., 2020), captured more phenotypic variance in Alzheimer’s disease than standard GWAS analysis methods (Lundberg et al., 2022). In addition, there is evidence that these findings replicated in an independent data set.

In conclusion, it is our expectation that RFlocalfdr with its direct and real-time capability of detecting trait-associated SNPs will greatly assist the analysis of genomic data.

## Software and data access

### R implementation

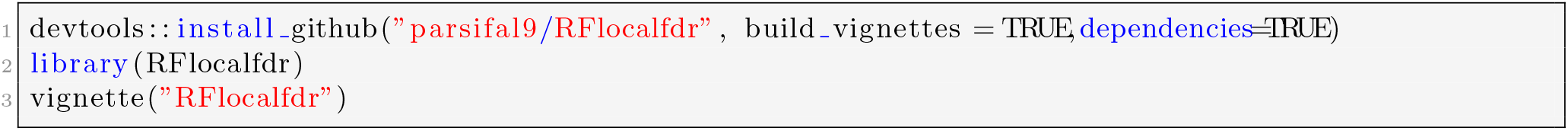

### Python implementation

Along with the R implementation of this tool, we released a python version included in the VariantSpark repository. The python script has been tested using python version 3.8.12. The libraries used are the following: numpy version 1.21.2 for numeric transformations; pandas version 1.4.1 to create and manage data frames; patsy version 0.5.2 to create the cubic regression splines; scipy version 1.7.3 to fit the data using the last squares method, compute the cumulative distribution functions and percentile point functions; and statsmodels version 0.13.2 to fit a generalized linear model.

Although the script contains several functions, the end user is only required to use the run_it_importances. This function requires a data frame as input with a single column containing the logarithm of the RF importance for each variant. Currently the method returns a dictionary with the FDR values, the estimates for the fitted function, and the p-values for the statistically significant variants.

Link to the python implementation: https://github.com/aehrc/VariantSpark/blob/3d9900e93f5a21768dfc0cf8733779ba147a3f48/python/varspark/pvalues_calculation.py (Notice that this implementation is the current last commit on a branch of VariantSpark)

## Supporting information

supplementary file 1 (Rmarkdown format)

supplementary file 2 (Rmarkdown format)

## Supplementary information

- supplementary_file_1.Rmd
- supplementary_file_2.Rmd

## Data Availability

- synthetic data – the code needed to generate it is provided.
- The 1000 Genome Project phase 3 dataset (Auton et al., 2015) was obtained from their FTP site as VCF files (section 3.2).
- The 64KX data set with 10,000 samples and 6 Million variables is taken from Bayat et al. (2020) where further information is given. It is based on the (section 3.3). was this data made public?

