## supplementary file 1 (Rmarkdown format) for "Thresholding Gini Variable Importance with a single trained Random Forest: An Empirical Bayes Approach": supplementary_file_1.Rmd

````
---
title: "Supplementary File 1 -- Simulated Data Example"
author: "Rob Dunne"
date: "Saturday, August  8, 2020"
output: 
    bookdown::html_document2:
      toc: true
      toc_depth: 2
      fig_caption: yes
vignette: >
  %\VignetteEngine{knitr::rmarkdown}
  %\VignetteIndexEntry{Supplementary File 1 -- Simulated Data Example}
  %\VignetteEncoding{UTF-8}
bibliography: paper.bib
header-includes:
  \usepackage{dcolumn}
---

```{r setup, include=FALSE}
knitr::opts_chunk$set(fig.width=6, fig.height=6, dpi=300,echo = FALSE)
knitr::opts_chunk$set(fig.pos = "H", out.extra = "")

```

# The data

We simulate a small data set with a covariance structure to explore the actions of **RFlocalfdr**.
This package implements an empirical Bayes method of determining a significance level for the Random Forest 
MDI (Mean Decrease in Impurity) variable importance measure. 
See @dunneThresholdingGiniVariable2022 for details

We 

- simulate the data set
- fit a Random Forest model
- use RFlocalfdr to estimate the significant variables
-  estimate the significant variables using

   - Boruta (@Kursa.and.Rudnicki.2010)
   - Recursive Fearure Elimination
   - AIR (@{Nembrini.et.al.2018})
   - PIMP (@Altmann.et.al.2010) 

and compare the results.


## setup
We plot a small data set of $300\times  508$ to explore the data generation method.
This dataset consist of _bands_, with _blocks_ of $\{1, 2, 4, 8, 16, 32, 64\}$ of
identical variables (see figure \@ref(fig:simulation2)). The variables are $\in \{0,1,2\}$, which is a common encoding for
genomic data where the numbers represent the number of copies of the minor allele. Only band 1 is used to calculate the
$y$ vector and $y$ is 1 if any of $X[, c(1, 2, 4, 8, 16, 32, 64)]$ is non-zero, so $y$ is unbalanced, containing more
1's than 0's.  We plot a small data set of $300\times  508$ to explore the data generation method.


```{r eval=FALSE,echo=TRUE}
library(ranger)
library(RFlocalfdr)
library(caret)
library(pROC)

```
```{r ,  eval=FALSE,echo=FALSE}
packageDescription("RFlocalfdr")$GithubSHA1

```
```{r eval=FALSE,echo=TRUE}
set.seed(13)
num_samples <- 300
num_bands <- 4
band_rank <- 6
num_vars <- num_bands * (2 ** (band_rank+1) -1)
print(num_vars)

X <- matrix(NA, num_samples , num_vars)
set.seed(123)

temp<-matrix(0,508,3)
var_index <- 1
for(band in 1:num_bands) {
     for (rank in 0:band_rank) {
         for (i in 1:2**rank) {
           temp[var_index,]<-c(band , rank, var_index)
             var_index <- var_index +1 
         }
     }
}

#png("./supp_figures/small_simulated_data_set.png")
plot(temp[,1],ylim=c(0,10),type="p") 
lines(temp[,2],type="b",col="red")
legend(0,10,c("band","rank"),pch=1,col=c(1,2))
#dev.off()
abline(v=c( 1,  2 , 4 , 8 ,16 ,32, 64 ))

table(temp[temp[,1]==1,2])
## # 0  1  2  3  4  5  6 
## # 1  2  4  8 16 32 64 

```

```{r simulation2, echo=FALSE, fig.cap="A representation of the simulation setup for the small data set. Each band contains blocks of size {1, 2, 4, 8, 16, 32, 64}, and each block consists of correlated (identical variables).", fig.align="center", out.width = '50%'}
knitr::include_graphics("./supp_figures/small_simulated_data_set.png")

```

## data
We have 50 bands and 300 observations so $X$ is $300 \times 6350$ with 127 non-null variables.  We fit a
standard RF to this dataset and record the resulting MDI importance score.

```{r eval=FALSE,echo=TRUE}
set.seed(13)
num_samples <- 300
num_bands <- 50
band_rank <- 6
num_vars <- num_bands * (2 ** (band_rank+1) -1)
print(num_vars)

X <- matrix(NA, num_samples , num_vars)
set.seed(123)
  
var_index <- 1
for(band in 1:num_bands) {
    for (rank in 0:band_rank) {
#        cat("rank=",rank,"\n")
        var <- sample(0:2, num_samples, replace=TRUE)
        for (i in 1:2**rank) {
            X[,var_index] <- var
            var_index <- var_index +1 
        }
#        print(X[1:2,1:140])
#        system("sleep 3")
    }
}

y <- as.numeric(X[,1] > 1 |  X[,2] > 1  |  X[,4] > 1 |  X[,8] > 1 |  X[,16] > 1 |
                X[,32] > 1 | X[,64] > 1 )

```

# Random forest
```{r eval=FALSE,echo=TRUE}
data <- cbind(y,X)
colnames(data) <- c("y",make.names(1:num_vars))
rfModel <- ranger(data=data,dependent.variable.name="y", importance="impurity",
             classification=TRUE,  mtry=100,num.trees = 10000, replace=TRUE, seed  = 17 )
imp <-log(ranger::importance(rfModel))
t2 <-count_variables(rfModel)
plot(density(imp))

library(pROC)
roccurve <- roc(c(rep(1,127),rep(0,6350-127)),imp)
plot(roccurve)
auc(roccurve) #0.9912

```


## Determine cutoff
```{r eval=FALSE,echo=TRUE}
cutoffs <- c(0,1,4,10)
#png("./supp_figures/simulated_data_determine_cutoff.png")
res.con<- determine_cutoff(imp,t2,cutoff=cutoffs,plot=c(0,1,4,10))
#dev.off()

``` 
We plot the kernel density estimate of the histogram of the data $y$, and the skew normal fit, $t_1$, using the data up to the
quantile $Q$, shown in red.

```{r , echo=FALSE, fig.cap="For this data set, the selected cutoff value is 0.", fig.align="center", out.width = '50%'}
knitr::include_graphics("./supp_figures/simulated_data_determine_cutoff.png")

```
```{r eval=FALSE,echo=TRUE}
plot(cutoffs,res.con[,3])
cutoffs[which.min(res.con[,3])]

```
by plotting $\max(|y - t_1|)$ versus the cutoff values, we determine the appropriate cutoff. In this case it is 
just $t2>0$


## Rflocalfdr
```{r eval=FALSE,echo=TRUE}
temp<-imp[t2 > 0]
#png("./supp_figures/simulated_data_plot_Q.png")
qq <- plotQ(temp,debug.flag = 1)
#dev.off()

```
Inspect the plot and see if the penalized selection procedure has succeeded.
In this case it has.


```{r , echo=FALSE, fig.cap="The Q value is selected.", fig.align="center", out.width = '50%'}
knitr::include_graphics("./supp_figures/simulated_data_plot_Q.png")

```

```{r eval=FALSE,echo=TRUE}

sensitivity<-function (data, positive = rownames(data)[1], ...) 
{
    numer <- sum(data[positive, positive])
    denom <- sum(data[, positive])
    sens <- ifelse(denom > 0, numer/denom, NA)
    sens
}

```
```{r eval=FALSE,echo=TRUE}
ppp<-run.it.importances(qq,temp,debug.flag = 0)
aa<-significant.genes(ppp,temp,cutoff=0.05,debug.flag=0,do.plot=2)
length(aa$probabilities) # 95

tt1 <-as.numeric(gsub("X([0-9]*)","\\1",names(aa$probabilities)))
tt2 <- table(ifelse((tt1 < 127),1,2))
# 1  2 
#59 36 
# The false discovery rate is 0.3789474
tt2[2]/(tt2[1]+tt2[2])
#59 36   36/(36+59) = 0.3789474

predicted_values<-rep(0, 6350);predicted_values[tt1]<-1

conf_matrix<-table(c(rep(1,127),rep(0,6350-127)),predicted_values)
conf_matrix
#   predicted_values
#       0    1
#  0 6187   36
#  1   68   59
#
#predicted_values
#       0    1
#  0   TN   FP
#  1   FN   TP

# sensitivity 0.464 TP/(TP+FN)
conf_matrix[2,2]/(conf_matrix[2,2] +conf_matrix[2,1])

#specificity 0.994 TN/(FP+TN)
conf_matrix[1,1]/(conf_matrix[1,2] +conf_matrix[1,1])

roccurve <- roc(c(rep(1,127),rep(0,6350-127)),predicted_values)
plot(roccurve)
auc(roccurve) #0.7294

accuracy<-(conf_matrix[1,1]+conf_matrix[2,2])/(sum(conf_matrix))
accuracy # 0.983622

```
The FDR is 0.379. We can also calculate the sensitivity, sensitivity and accruacy.


# Boruta

We try the Boruta algorithm (@Kursa.and.Rudnicki.2010).

```{r eval=FALSE,echo=TRUE}
library(Boruta)
set.seed(120)  
boruta1 <- Boruta(X,as.factor(y),  num.threads = 7,getImp=getImpRfGini)
print(boruta1)
#Boruta performed 99 iterations in 19.54727 secs.
#4 attributes confirmed important: X4859, X58, X6132, X7;
# 6346 attributes confirmed unimportant: X1, X10, X100, X1000, X1001 and 6341 more;
plotImpHistory(boruta1)
aa <- which(boruta1$finalDecision=="Confirmed") 
bb <- which(boruta1$finalDecision=="Tentative") 
predicted_values <-rep(0,6350);predicted_values[c(aa,bb)]<-1
conf_matrix<-table(c(rep(1,127),rep(0,6350-127)),predicted_values)
conf_matrix
#   predicted_values
#       0    1
#  0 6221    2
#  1  125    2

# sensitivity 0.016 TP/(TP+FN)
conf_matrix[2,2]/(conf_matrix[2,2] +conf_matrix[2,1])

#specificity 0.999 TN/(FP+TN)
conf_matrix[1,1]/(conf_matrix[1,2] +conf_matrix[1,1])

roccurve <- roc(c(rep(1,127),rep(0,6350-127)),predicted_values)
plot(roccurve)
auc(roccurve) #0.5077

accuracy<-(conf_matrix[1,1]+conf_matrix[2,2])/(sum(conf_matrix))
accuracy #0.98

```
```{verbatim, eval=FALSE,echo=TRUE}
  D  !D
+ TP FP
- FN TN
```

- Sensitivity=[TP/(TP+FN)]×100
- Specificity=[TN/(FP+TN)]×100
- Positive predictive value(PPV)= [TP/(TP+FP)]×100
- Negative predictive value(NPV)=[TN/(FN+TN)]×100.

# Recursive Fearure Elimination
This is provided by the package Pomona (@Pomona.2022)
```{r eval=FALSE,echo=TRUE}
library(Pomona)
set.seed(4)
colnames(X) <- make.names(1:dim(X)[2])
system.time(res <- var.sel.rfe(X, y, prop.rm = 0.2,  recalculate = TRUE, tol = 10, 
    ntree = 500, mtry.prop = 0.2, nodesize.prop = 0.1, no.threads = 7, 
    method = "ranger", type = "classification", importance = "impurity", 
    case.weights = NULL) )
res$var


for ( ii in 1:500){
set.seed(ii)
colnames(X) <- make.names(1:dim(X)[2])
system.time(res <- var.sel.rfe(X, y, prop.rm = 0.2,  recalculate = TRUE, tol = 10, 
    ntree = 500, mtry.prop = 0.2, nodesize.prop = 0.1, no.threads = 7, 
    method = "ranger", type = "classification", importance = "impurity", 
    case.weights = NULL) )
#res$var
cat(ii, " ",length(intersect(res$var,aa)),"\n")
}

# [1] "X1"    "X108"  "X12"   "X13"   "X14"   "X18"   "X19"   "X2"    "X25"  
#[10] "X29"   "X3"    "X4"    "X5"    "X50"   "X53"   "X5469" "X6"    "X61"  
#[19] "X6135" "X62"   "X65"   "X7"    "X72"   "X81"  
,
aa<-c("X1", "X108",  "X12",   "X13"  , "X14" ,  "X18"  , "X19"  , "X2", "X25" , 
 "X29" ,  "X3", "X4", "X5", "X50" ,  "X53" ,  "X5469", "X6", "X61"  ,
 "X6135", "X62" ,  "X65"  , "X7", "X72" ,  "X81"  )

tt<-as.numeric(gsub("X([0-9]*)","\\1", res$var))
table(ifelse((tt < 127),1,2))
# 1  2 
#22  2 0.0833

res<-c(1, 108,12, 13, 14, 18, 19, 2,25,29, 3,4,5,50, 53, 5469,6,61, 6135,62, 65, 7,72, 81)
predicted_values <-rep(0,6350);predicted_values[c(res)]<-1
conf_matrix<-table(c(rep(1,127),rep(0,6350-127)),predicted_values)
conf_matrix
#   predicted_values
#       0    1
#  0 6221    2
#  1  105   22

# sensitivity 0.1732283 TP/(TP+FN)
conf_matrix[2,2]/(conf_matrix[2,2] +conf_matrix[2,1])

#specificity 0.9996  TN/(FP+TN)
conf_matrix[1,1]/(conf_matrix[1,2] +conf_matrix[1,1])

roccurve <- roc(c(rep(1,127),rep(0,6350-127)),predicted_values)
plot(roccurve)
auc(roccurve) #0.5865
accuracy<-(conf_matrix[1,1]+conf_matrix[2,2])/(sum(conf_matrix))
accuracy #  0.9831496

```

# AIR

See @{Nembrini.et.al.2018}. AIR is provided in the package _ranger_, using the option _impurity_corrected_.
```{r eval=FALSE,echo=TRUE}
rfModel2 <- ranger(data=data,dependent.variable.name="y", importance="impurity_corrected",
       classification=TRUE,  mtry=100,num.trees = 10000, replace=TRUE, seed  = 17 )

ww <- importance_pvalues( rfModel2, method = "janitza")
p <- ww[,2]
cc <- which(p< 0.05)
tt <- as.numeric(gsub("X([0-9]*)","\\1", cc))
tt2 <- table(ifelse((tt < 127),1,2))
#  1   2 
#127 273    0.6825
predicted_values <-rep(0,6350);predicted_values[tt]<-1
conf_matrix<- table(c(rep(1,127),rep(0,6350-127)),predicted_values)
conf_matrix
#>    predicted_values
#       0    1
#  0 5950  273
#  1    0  127

# sensitivity 1 TP/(TP+FN)
conf_matrix[2,2]/(conf_matrix[2,2] +conf_matrix[2,1])

#specificity 0.95613 TN/(FP+TN)
conf_matrix[1,1]/(conf_matrix[1,2] +conf_matrix[1,1])


roccurve <- roc(c(rep(1,127),rep(0,6350-127)),predicted_values)
plot(roccurve)
auc(roccurve) # 0.9781
accuracy<-(conf_matrix[1,1]+conf_matrix[2,2])/(sum(conf_matrix))
accuracy #  0.9570079

```

# PIMP

We take the @{vita.2015} _Variable Importance Testing Approaches_ 

```{r eval=FALSE,echo=TRUE}
library(vita) #vita: Variable Importance Testing Approaches
source("/home/dun280/Dropbox/R_libraries/RF_localfdr/RFlocalfdr/R/my.pimp.s")
y<-factor(y)
X<-data.frame(X)
cl.ranger <- ranger(y~. , X,mtry = 3,num.trees = 1000, importance = 'impurity')
system.time(pimp.varImp.cl<-my.ranger.PIMP(X,y,cl.ranger,S=10, parallel=TRUE, ncores=3)) 
pimp.t.cl <- PimpTest(pimp.varImp.cl,para = FALSE)
aa <- summary(pimp.t.cl,pless = 0.1)

predicted_values <-rep(0,6350);predicted_values[which(aa$cmat2[,"p-value"]< 0.05)]<-1
conf_matrix<-table(c(rep(1,127),rep(0,6350-127)),predicted_values)
conf_matrix
#       0    1
#  0 5678  545
#  1   88   39

#sensitivity 0.307
conf_matrix[2,2]/(conf_matrix[2,2] +conf_matrix[2,1])

#specificity 0.9124 TN/(FP+TN)
conf_matrix[1,1]/(conf_matrix[1,2] +conf_matrix[1,1])

roccurve <- roc(c(rep(1,127),rep(0,6350-127)),predicted_values)
plot(roccurve)
auc(roccurve) #  0.6089
accuracy<-(conf_matrix[1,1]+conf_matrix[2,2])/(sum(conf_matrix))
accuracy # 0.8985827


tt<-as.numeric(gsub("X([0-9]*)","\\1",  names(which(aa$cmat2[,"p-value"]< 0.05))))
table(ifelse((tt < 127),1,2))
# 1  2 
# 39 556      556 /(556+39) = 0.9344538


```{r eval=FALSE,echo=FALSE}
library(tables)
X <- rnorm(125, sd=100)
Group <- factor(sample(letters[1:5], 125, rep=TRUE))
tab <- tabular( Group ~ (N=1)+Format(digits=2)*X*((Mean=mean) + Heading("Std Dev")*sd) )
table_options(knit_print = FALSE)
tab        # This chunk uses the default results = 'markup'

toHTML(tab)  # This chunk uses results = 'asis'
table_options(CSS =
"<style>
#ID .center { 
  text-align:center;
  background-color: aliceblue;
}
</style>", doCSS = TRUE)
tab
table_options(doCSS = FALSE)
```


# Conclusion


```{r table3, echo=FALSE, message=FALSE, warnings=FALSE, results='asis'}

tabl <- "
    Method     | true positives | false positives | Sensitivity        | AUC            |
    RFlocalfdr | 59             | 36              | 0.99               | 0.73           |
    Boruta     | 2              | 2               | 0.99               | 0.50           |
    RFE        | 22             | 2               | 0.99}              | 0.58           |
    AIR        | 176            | 273             | 0.96               | 0.98           |
    PIMP       | 39             | 556             | 0.91               | 0.61           |
--------------------------------------------------------------------------- 
"
cat(tabl) # output the table in a format good for HTML/PDF/docx conversion
```


Performance of variable selection for the simulated example by several methods. The best outcomes for each category are in bold face.
RFE has performed better than Boruta but apart from that, ordering the methods will depend on the cost of false positives versus the cost of false negatives.


How do I get boldface in a table? neither  **0.99**  or  $\textbf{0.99}$ work


# References

<!-- tabular( -->
<!-- \begin{table}[tbhp] -->
<!--   \centering -->
<!--   \begin{tabular}{||p{2cm}|l|l|l|l|l|l|l||} -->
<!--     \hline -->
<!--     Method     & true positives & false positives & Sensitivity & AUC  \\ -->
<!--     \hline -->
<!--     RFlocalfdr & 59             & 36              & \textbf{0.99}       & 0.73 \\ -->
<!--     Boruta     & 2              & 2               & \textbf{0.99}        & 0.50 \\ -->
<!--     RFE        & 22             & 2               & \textbf{0.99}        & 0.58 \\ -->
<!--     AIR        & 176            & 273             & 0.96        & \textbf{0.98} \\ -->
<!--     PIMP       & 39             & 556             & 0.91        & 0.61 \\ -->
<!--     \hline -->
<!--   \end{tabular} -->
<!--   \caption{Performance of variable selection for the simulated example by several methods. The best outcomes for each -->
<!--     category are in bold face. -->
<!--   %RFE has performed better than Boruta but apart from that, ordering the methods will depend on the cost of false positives versus the cost of false negatives. -->
<!--   } -->
<!--   \label{table:simulation_results} -->
<!-- \end{table} -->
<!-- ) -->

<!--  rmarkdown::html_vignette   - figure references do not work, get figure \@ref(fig:simulation2) -->
<!--  bookdown::html_document2:  -  figure references  work, get figure 1     -->
````
