## supplementary file 2 (Rmarkdown format) for "Thresholding Gini Variable Importance with a single trained Random Forest: An Empirical Bayes Approach": supplementary_file_2.Rmd

````
---
title: "Supplementary File 2 -- 1000 Genome Data"
author: "Rob Dunne"
date: "Saturday, August  8, 2020"
output:	 
  bookdown::pdf_document2:
    toc: true
    toc_depth: 2
    fig_caption: yes
header-includes:
  - \usepackage{placeins}
  - \usepackage{float}
  - \usepackage{siunitx}
  - \usepackage{dcolumn}
bibliography: paper.bib

---


```{r setup, include=FALSE}
knitr::opts_chunk$set(fig.width=6, fig.height=6, dpi=300,echo = FALSE)
knitr::opts_chunk$set(fig.pos = "H", out.extra = "")

```

```{r eval=FALSE,echo=FALSE}
rm(list=ls())
library(RFlocalfdr)
packageDescription("RFlocalfdr")$RemoteRef #[1] #
#library(goftest)
#library(diptest)
#source("~/Dropbox/random_forests/exploration/script19/scripts.s") what was this for?
library(plotrix)

```

# Introduction
To demonstrate the applicability of RFlocalfdr on a real-world dataset, we applied it to the 1000 Genome
Project phase 3 dataset (@autonGlobalReferenceHuman2015). 
We show how to download and pre-process the data, and fit a RF model.
We provide the outputs of the RF model (the variable importances and counts) with the **RFlocalfdr** package.


# The Data
We obtained the 1000 Genomes Project data as
VCF files from their FTP site. Each VCF file contains the SNP variants for every individual across one
chromosome. There are 2,504 individuals and we limit the analysis to the 1M SNPs on chromosome 22. We
train a RF to predict the ethnicity of each individual.

# Processing
Some pre-processing is necessary to get the data into a format for use in R.

## get the data 

```{bash eval=FALSE,echo=TRUE}
> wget  http://hgdownload.cse.ucsc.edu/gbdb/hg19/1000Genomes/phase3/ALL.chr22.phase3_shapeit2_mvncall_integrated_v5a.20130502.genotypes.vcf
> wc ALL.chr22.phase3_shapeit2_mvncall_integrated_v5a.20130502.genotypes.vcf 
#1103800
> tail -1103547 ALL.chr22.phase3_shapeit2_mvncall_integrated_v5a.20130502.genotypes.vcf | \
    cut --complement -f 1-9 |  \
	sed -e 's/0|0/0/g'  -e 's/0|[1-9]/1/g' -e 's/[1-9]|0/1/g'  -e 's/[1-9]|[1-9]/2/g' > tt3

# set up the processing environment
> salloc --mem=1000GB --nodes=1  --ntasks-per-node 64 -J interactive -t 5:00:00 srun --pty /bin/bash -l

```
## transpose the data
We run a batch file to transpose the data

- data in ALL.chr22....vcf is $p \times n$ (253 line header)
- data in **tt3** is $p \times n$
- data in **ch22.csv** is $n \times p$, saved as **aa2.Rdata**, 121M


```{r eval=FALSE,echo=TRUE}
#R CMD BATCH --no-save  --no-restore --no-readline transpose.s trainspose.s.out &
library(data.table)
system.time(aa2<-fread("tt3",sep="\t",nrows=1103547,skip=0))
system.time(aa3<-t(aa2))
system.time(aa3<-as.data.frame(aa3))
system.time(fwrite(aa3, "ch22.csv", sep = "\t", col.names = FALSE))

```
## prepare the data
```{r eval=FALSE,echo=TRUE}
y<-read.table("Y.csv")
library(data.table)
system.time(aa2<-fread("ch22.csv",sep="\t",nrows=2504,skip=0))
#Read 2504 rows and 1103548 (of 1103548) columns from 5.147 GB file in 00:11:54
#   user  system elapsed 
#771.320   4.728 775.542 

#dim(aa2) #2504 1103548
#system.time(aa2<-as.data.frame(aa2))

dim(aa2) # 2504 1103547
system.time(aa2<-data.frame(y,aa2))
aa[1:10,1:10]
save(aa2,file="aa2.Rdata")

table(as.numeric(factor(aa2[,1])),aa2[,1])
#    AFR AMR EAS EUR SAS
#  1 661   0   0   0   0
#  2   0 347   0   0   0
#  3   0   0 504   0   0
#  4   0   0   0 503   0
#  5   0   0   0   0 489

```

## fit the RF model
Now that we have the data in the correct format we can fit a RF model.

```{r eval=FALSE,echo=TRUE}
system.time(fit.ranger.7 <- ranger(dependent.variable.name= "V1", data = aa2,
                                 importance = "impurity",
                                  num.threads=20,num.trees = 100000,
                                 seed=123))
#   user      system   elapsed 
# 56614.757    50.264  3565.236
## Ranger result
## Call:
## ranger(dependent.variable.name = "V1", data = aa2, importance = "impurity", 
##                               num.threads = 20, num.trees = 1e+05, seed = 123) 
## Type:                             Classification 
## Number of trees:                  1e+05 
## Sample size:                      2504 
## Number of independent variables:  1103547 
## Mtry:                             1050 
## Target node size:                 1 
## Variable importance mode:         impurity 
## Splitrule:                        gini 
## OOB prediction error:             4.27 %

```

<!-- refers to a fitted model in  pearcey:/home/dun280/thousand_genomes/chr22_phase3_dpas03/script4.s -->
<!-- imp7 -- importances   #1103547 -->
<!-- t2   --  counts -->
<!-- can we get names for imp7 and start t2 at 1 not 0 -->
<!-- we have an imp1 in the package -- need better names, also why is it loaded automaticaly? -->

# RFlocalfdr
## starting from the saved importance values
We provide the outputs of the RF model (the variable importances and counts) with the **RFlocalfdr** package.
Figure \@ref(fig:importances) shows a density plot of the log importances. The multi-modality of the distribution is the first problem.
The target density, which is where we will apply the empirial Bayes argument, is outlined in red.

```{r eval=FALSE,echo=TRUE}
library(RFlocalfdr)
data(ch22)
imp <- ch22[[1]]
t2 <- ch22[[2]]

```

```{r eval=FALSE,echo=TRUE}
#png("./supp_figures/density_imp7.png")
plot(density(log(imp)),xlab="",ylab="Density",axes=FALSE,cex.lab=1.4,cex.main=1.5,main="log importances may be multi-modal")
draw.ellipse(c(-5.4), c(0.07), c(1.5), c(0.05), col=NA, angle=c(0),border="red",lwd=4)
box()
#dev.off()
imp1<-log(imp)
length(imp1[imp1 != -Inf]) #629181

```

```{r importances, echo=FALSE, fig.cap="The importances may be multi-model.", fig.align="center", out.width = '50%'}
knitr::include_graphics("./supp_figures/density_imp7.png")

``` 

## Selecting the cutoff $C$

- count the number of times that each variable is used -- this is $C$
- progressively threshold $C$ giving  $f_c (z) = f (z) | C> c$
-  for $f_c (z)$, we can fit a Skew-Normal distribution (@Azzalini.2022)
-Our current procedure is

   - for the distribution $f_c (z)$, fit a  Skew-Normal up to the $q^{th}$ quantile, $S_q(z)$
   - calculate the $L_\infty$  norm $d_c = \max_z | f_c (z) - S_q(z)|$

- plot $d_c$ versus $c$ (figure \@ref(fig:determinecutoff)) and pick the minimum value $c^\star$. Call the
  corresponding distribution $f_{c^\star} (z)$
- we shift  $f_{c^\star} (z)$ along the $z$ positive axis so the smallest value is 0. 

This removed 87\% of the variables and results in the density shown in figure \@ref(fig:thousandgenomesRFlocfdr).


```{r eval=FALSE,echo=TRUE}

cutoffs <- c(0,1,2,3,4,5,6,10,11,13,15,16,17,19,20,30,40,50)
res.con<- determine_cutoff(imp1,t2,cutoff=cutoffs)
cutoffs[which.min(res.con[,3])]

#png("./supp_figures/determine_cutoff.png")
plot(cutoffs,res.con[,3],main="determine cutoff")
abline(v=30,col="blue")
#dev.off()

temp<-imp1[t2 > 30]   #
temp<-temp[temp != -Inf]
temp <- temp - min(temp) + .Machine$double.eps

#png("./supp_figures/determine_cutoff_30.png")
plot(density(temp),main="cutoff=30")
#dev.off()

```

```{r determinecutoff, echo=FALSE, fig.align="center",fig.cap="determine cutoff", out.width = '50%'}
knitr::include_graphics("./supp_figures/determine_cutoff.png")

``` 


## RFlocalfdr 

RFlocalfdr returned 6650 SNPs  as significant at a FDR of 0.2.
See Figure \@ref(fig:thousandgenomesRFlocfdr) for the diagnostic plot. 


```{r eval=FALSE,echo=TRUE}
qq <- plotQ(temp,debug.flag = 1)

ppp<-run.it.importances(qq,temp,debug=0)

#png("./supp_figures/thousand_genomes_RFlocfdr.png")
aa<-significant.genes(ppp,temp,cutoff=0.2,do.plot=2)
#dev.off()

length(aa$probabilities)
#[1] 6650

```

```{r eval=FALSE,echo=FALSE}
RFlocalfdr.bb<-significant.genes(ppp,temp,cutoff=0.05,do.plot=2)
length(RFlocalfdr.bb$probabilities)
#3653

ggvenn(list(RFlocalfdr_0.2= names(aa$probabilities),RFlocalfdr_0.05= names(RFlocalfdr.bb$probabilities)))
aa.eb<-names(aa$probabilities)
aa.eb<- as.numeric(gsub("V([0-9]*)","\\1",aa.eb))

```


```{r thousandgenomesRFlocfdr, echo=FALSE, fig.align="center",fig.cap="Diagnostic plot output from RFlocalfdr", out.width = '50%'}
knitr::include_graphics("./supp_figures/thousand_genomes_RFlocfdr.png")

``` 


# Boruta

The Boruta algorithm (@Kursa.and.Rudnicki.2010) returned a similar number (6,773) of significant SNPs (see figure
\@ref(fig:venndiagramsiggenes). Unlike RFlocalfdr, which offers a continuous $p$-value scale for the adjustment
of FDR, Boruta offers only two levels, and returns 1,443 at the more conservative "Confirmed" option compared to the
"Tentative" option. All of the ``Confirmed'' SNPs are in our 6,335 list and the overlap of our set and the combined
"Confirmed" and "Tentative" is 0.82 (see Figure \@ref(fig:venndiagramsiggenes)). The "Confirmed" SNP have
RFlocalfdr $p$-values that cluster at the low end. A one-way ANOVA test testing for association of RFlocalfdr $p$-values
and Boruta categories give a $p$-value $< 2\times 10^{-16}$.

<!-- see file petrichor://home/dun280/workspace/thousand_genomes/chr22_phase3_dpas03/script4.s -->
<!-- > locate boruta_variables_Confirmed.Rdata -->
<!-- /home/dun280/Dropbox/random_forests/exploration/script19/boruta_variables_Confirmed.Rdata -->
<!-- > grep  boruta_variables_Confirmed.Rdata * -->
<!-- > temp1.Rnw:load("./script19/boruta_variables_Confirmed.Rdata") -->
<!-- > temp.Rnw:load("./script19/boruta_variables_Confirmed.Rdata") -->
<!-- > t.Rnw:load("./script19/boruta_variables_Confirmed.Rdata") -->


```{r eval=FALSE,echo=TRUE} 
library("Boruta")
system.time(boruta2 <- Boruta(X,y,getImp=getImpRfGini,num.threads=40))
#     user    system   elapsed 
#36068.385  2320.502 12496.694

plotImpHistory(boruta2)
getConfirmedFormula(boruta2)
boruta.Confirmed <- which(boruta2$finalDecision=="Confirmed")
boruta.Tentative <-which(boruta2$finalDecision=="Tentative")

length(intersect(aa.eb,boruta.Tentative)) #[1] ]  4029
length(intersect(aa.eb,c(boruta.Confirmed,boruta.Tentative))) #5472

```


# PIMP
<!-- see file petrichor://home/dun280/workspace/thousand_genomes/chr22_phase3_dpas03/script3.s -->

PIMP (@Altmann.et.al.2010) is provided by the **vita** package (@vita.2015). The PIMP algorithm is described in
@Altmann.et.al.2010 for permutation importance.  The **my.PIMP** function provided in this package is based on the
**vita** implementation but has been modified so that:

- it can be applied to MDI importance
- it accepts the output of a **ranger** random forest as well as the **randomForest** package
- my.pimp.s is not in the RFlocalfdr package

PIMP preforms a test on each variable and so can be corrected for multiple testing. This reduces the number of 
significant variables from 787502 to 709270, not a huge reduction.

```{r eval=FALSE,echo=TRUE}
library(ranger)
library(vita) #vita: Variable Importance Testing Approaches
source("/home/dun280/Dropbox/R_libraries/RF_localfdr/RFlocalfdr/R/my.pimp.s") 
system.time(load("aa2.Rdata"))
X<-aa2[,-1]
y<-aa2[,1]
system.time(pimp.varImp.cl<-my.ranger.PIMP(X,y,fit.ranger.7,S=100, parallel=TRUE, ncores=32))


save(pimp.varImp.cl,file="./script3/pimp.varImp.cl.Rdata")
load(file="~/Dropbox/random_forests/exploration/script3/pimp.varImp.cl.Rdata")
pimp.t.cl <- PimpTest(pimp.varImp.cl,para = FALSE)
aa <- summary(pimp.t.cl,pless = 0.1)
class(aa) #[1] "summary.PimpTest"

pvalue  <- aa$cmat2[,"p-value"]
length(which(pvalue< 0.05) ) #787502

library(fdrtool)
fdr.out <- fdrtool(pvalue, statistic = "pvalue", plot = TRUE, verbose = TRUE, cutoff.method = "fndr", color.figure = TRUE)
sum(fdr.out$lfdr < 0.05) #[1] 709270


pimp.varImp.cl
> system.time(cl.ranger <- ranger(y=y ,x= X,num.trees = 500, importance = 'impurity',num.threads=128))
   user  system elapsed
626.192   8.940  75.117
> system.time(pimp.varImp.cl<-my.ranger.PIMP(X,y,cl.ranger,S=100, parallel=TRUE, ncores=64))
     user    system   elapsed
22503.670  7070.980   976.964


```

# AIR

-  so how does the AIR paper do it?

<!-- see script3 on petra -- impurity corrected -->

- note the $p$-value has been set to 0.000000001 
- why and how to justify?

```{r eval=FALSE,echo=TRUE}
system.time(fit.ranger.3 <- ranger(dependent.variable.name= "V1", data = aa2,
                                 importance = "impurity_corrected",
                                 num.threads=50,num.trees = 100000,
                                 seed=123))
#     user     system    elapsed 
#125322.943    256.062   2976.523
#OOB prediction error:             5.03 %

ww<- importance_pvalues( fit.ranger.3, method = "janitza")
p<-ww[,2]
cc<-which(p< 0.000000001)  #61092
cc<-which(p< 0.05)  #191489

```

```{r eval=FALSE,echo=FALSE}
save(cc,file="impurity_corrected_variables.Rdata")
load("./script19/impurity_corrected_variables.Rdata") #]  61092
length(intersect(names(aa$probabilities),names(cc)))  # 6650
library(ggvenn)  
#length(intersect(boruta.Tentative,cc))  #2781
#length(intersect(cc,c(boruta.Confirmed))) #1081

names.boruta.all <- paste("V",c(boruta.Confirmed,boruta.Tentative),sep="")
names.boruta.Confirmed <- paste("V",c(boruta.Confirmed),sep="")

ggvenn(list(RFlocalfdr= names(aa$probabilities),AIR= names(cc), Boruta =names.boruta.all),
       show_percentage = FALSE)
ggvenn(list(RFlocalfdr= names(bb$probabilities),AIR= names(cc), Boruta =names.boruta.Confirmed),
       show_percentage = FALSE)

ggvenn(list(RFlocalfdr= names(RFlocalfdr.bb$probabilities),AIR= names(cc), Boruta =names.boruta.Confirmed,
            RFE=res$var),
       show_percentage = FALSE)


library(venn)
x<-list(RFlocalfdr= names(RFlocalfdr.bb$probabilities),AIR= names(cc),
        Boruta =names.boruta.Confirmed,   PIMP=bb,  RFE=res$var)
#png("./script19/ven_diagram_sig_genes.png")
venn(x,zcolor = "style")
#dev.off()

```

## AIR + locfdr

- The distribution of importance values is designed to be symmetric about 0 (@Nembrini.et.al.2018). This opens the possibility of using the empirical Bayes approach 
of @Efron.2010a directly on the importances returned by AIR.

- This appeared to work well for the simulated data set (see supplementary_file_1), however we see problem for the 100 Genome data set
- there are 61092 variables with $p$-values of 0, so we will recover the same variables as AIR (see above)
- the distribution is highly spiked. It is not well modelled as a Normal or $t$ due to the number of extreme outliers
  


```{r eval=FALSE,echo=TRUE}
sum(ww[,"pvalue"]==0)  #61092
imp<- ww[,"importance"]

#png("AIR_locfdr_imp.png")
plot(density(imp),xlab="importances, impurity corrected",main="")
#dev.off()

temp<- ww[,"importance"][ww[,"importance"]  < 0.0001 ]
temp<- temp[temp > -0.0001 ]
length(temp) #[1] 818071
#png("AIR_locfdr_imp_reduced.png")
plot(density(temp),xlab="importances, impurity corrected",main="")
#dev.off()


mean.temp <- mean(temp)
var.temp <- var(temp)
temp <- (temp-mean.temp)/sqrt(var.temp)
library(locfdr)

#png("AIR_locfdr_plot.png")
aa<-locfdr(temp,mlests =  c(-0.117980663840164, 0.000225216379326981),df=20)
#dev.off()

sum(aa$fdr< 0.05)
#9980919

```


```{r AIRlocfdr, echo=FALSE, fig.align="center",fig.cap="The importance values returned by AIR, and the output plot from locfdr", out.width = '40%',fig.show="hold"}
knitr::include_graphics("./supp_figures/AIR_locfdr_imp.png")
knitr::include_graphics("./supp_figures/AIR_locfdr_plot.png")

``` 


# RFE

RFE returned 59 nested sets and selects the best set using the smallest prediction error. RFE appears to have
focused on specificity and only returns 12 SNPs, which were also found by all other methods (see Figure
\@ref(fig:venndiagramsiggenes)).

- this only used 500 trees

```{r eval=FALSE,echo=TRUE}
library(Pomona)
system.time(res <- var.sel.rfe(x=x, y=y, prop.rm = 0.2,
                   recalculate = TRUE, tol = 10, 
    ntree = 500, mtry.prop = 0.2, nodesize.prop = 0.1, no.threads = 40, 
    method = "ranger", type = "classification", importance = "impurity", 
    case.weights = NULL) )
# user       system  elapsed
#216804.28    320.18   7001.98

names(res) #[ 1] "info"      "var"       "info.runs"

res$var
# [1] "V1073032" "V128376"  "V212586"  "V585417"  "V640647"  "V690597" 
# [7] "V715130"  "V749345"  "V781170"  "V781330"  "V881168"  "V933772" 

plot(sort(match(res$var,names(bb$probabilities))))

```
# Runtime

It is easy to measure the run-time of a process but much harder to produce a meaningful comparison of techniques.
Issues that should be considered include

- will the code run in parallel? I have given both **user** and **elapsed** time and the number of cores
- what options should the code be called with? 

    - I am not running **ranger** for **RFlocalfdr** with the default settings. In this case we want to estimate the
  null distribution so we need sufficient trees so that the 1M variables have multiple chances of being considered at a
  node.  Setting this variable in an optimal way is related to the degree of correlation between the predictor
  variables. At the moment we simple set it to a very large value.
    - However it is not clear how we should now set the parameters for the other techniques. Setting the number of trees very high
	for methods that do not benefit from this may produce a misleading comparison. See table \ref{thousand_genome_results} for the 
	run times and parameters of the methods used in this document.
	
- should waiting time be included in the run-time. These results were obtained on a large shared cluster. I have not
reported waiting time except in the case of PIMP inf table \ref{systematic_thousand_genome_results} where the waiting
time appeared to be indefinite.  A notable feature of shared computing clusters with scheduling systems (such as
**slurm**) is that, while it is easy to request large amounts of resources, one may find that the waiting time for the job to start becomes
extremely long.

Table \ref{systematic_thousand_genome_results}

- the resources we set to 40 cores, 200GB of RAM and 5 hours run-time
- for Boruta and 100000 trees, increasing the time significantly (96 hours) did not help
- for PIMP, we progressively increased the memory until the scheduler seem to postpone the job indefinitely 


\begin{table}[tbhp]
  \centering
  \begin{tabular}{||l|l|D{.}{.}{3}|D{.}{.}{3}|D{.}{.}{3}|D{.}{.}{3}|l||}
    \hline
    Method     &  SNPs returned         &  \multicolumn{1}{c|}{elapsed}    &\multicolumn{1}{c|}{user}     &\multicolumn{1}{c|}{threads}   &\multicolumn{1}{c|}{trees}  &  evaluations/time     \\
    \hline
    RFlocalfdr & 6,335          &   3565.24  &  56614.76       &  20   &  100000 & 1     \\
    Boruta     & 6,773          & 12496.69   &  36068.38       & 40    &   500   & 100   \\
    RFE        & 12             &  7001.98   &  216804.28           &   40  & 500& 57      \\
    AIR        & 61,092         &  2976.52   & 125322.94     &  50 &  100000     & 1.5 -- 2       \\
    PIMP       & 787502         &  1052.08   &  23129.86           &    64 &  500& 100     \\
    \hline
  \end{tabular}  
  \caption{The elapsed and run times, threads and trees for the methods considered in this document.
  "evaluations/time" refers to the number of evaluations of the RF or, in the case of AIR, to the run-time. AIR evaluates the RF model once but with twice the number of
  variables. The "evaluations/time" for RFE is 57 but each run has a reduced number of variables are so will be faster.}
  \label{thousand_genome_results}
\end{table}


\begin{table}[tbhp]
  \centering
  \begin{tabular}{||l|l|D{.}{.}{3}|D{.}{.}{3}|D{.}{.}{3}|D{.}{.}{3}|c||}
    \hline
    Method     &  ntree        &\multicolumn{1}{c|}{SNPSs returned }   &\multicolumn{1}{c|}{elapsed}    &\multicolumn{1}{c|}{user} &  \multicolumn{1}{c|}{MaxRSS (GB)}   &notes           \\
    \hline
    RFlocalfdr &100000   &  6335          &   2025.17   &  66677.73     &  79.40       &               \\
  RFlocalfdr & 10000&   3176         &    274.041   &    6779.205     &   76.11       &               \\
   RFlocalfdr &  1000 & 580         &  92.73     & 739.057     &  76.11        &         \\
Boruta     & 100000  &  -             &  -          &  -            &              &  out of time  \\
  Boruta     & 10000& -             &  -          &  -            &   -     &  out of time  \\
    Boruta     & 1000 &  12081       & 12442.77   & 67078.45    &  66.21   &   \\
RFE        & 100000  &  -             &  -           &   -           &   -         &  out of time  \\
  RFE        &  10000& -             &  -           &   -          &   -         &  out of time  \\
    RFE        & 1000 &  -            &    -        &   -          &     -     &   out of time\\
AIR        & 100000  & 191489        &  2976.52    & 125322.94     &  77.72      &               \\
   AIR        &  10000&155154        &  2976.52    & 9940.345       &  76.13      &               \\
      AIR        & 1000 & 88225        &  107.88    &1114.116     &  76.11   &    \\
PIMP       & 100000  &  -            &     -       &    -          &       -      & out of memory    \\
    PIMP       &  10000& -            &     -       &    -          &       -      & out of memory    \\
    PIMP       &  1000 &  -          &    -         &  -           & -         & out of memory  \\
\hline
  \end{tabular}
     \caption{
    A more systematic, but still somewhat arbitrary, attempt to measure the run-times. The files for replicating this
   trial are included in the **RFlocalfdr** package.
   }
    \label{systematic_thousand_genome_results}
\end{table}


# Conclusion

RFlocalfdr offers the option to adjust the sensitivity/specificity trade-off, by adjusting the cutoff $p$-value of the
selected SNPS. Likewise for RFE, we could select another of the nested sets with a larger or smaller number of
elements.  However Boruta and AIR do not offer this flexibility as there is no criteria for sub-setting the set of selected
SNPs.

As in the simulated example, AIR (@Nembrini.et.al.2018) selects an order of magnitude more significant SNPs
(61,092) than RFlocalfdr and Boruta, potentially having a high sensitivity.  However, with the $p$-value of all 61,092
SNPs receiving a score 0, AIR does not offer any option for tuning the specificity and the resulting false positives
rate may be substantial (see Figure \@ref(fig:venndiagramsiggenes)).


AIR is  very much the outlier with a vastly larger number of selected SNPS. Whether this is desirable or not would depend
on the subsequent analysis, which might involve building a classifier, understanding functional pathways, or doing
further laboratory work on selected SNPS.

We next evaluated the run-time. Since RFlocalfdr does not require multiple passes of the RF to be conducted for the
statistical analysis, the run-time is essentially the run-time for the RF model. This can be run in parallel so the
speed will depend on the number of cores. Using the same processing parameters, we found that RFlocalfdr was 25 times
faster than the next best performing method, AIR, 57 times faster than RFE and 100 times faster than Boruta.

A Venn diagram of the overlaps in SNP for RFlocalfdr, Boruta, AIR, RFE and PIMP. AIR and PIMP are very much the
outliers, with more than $10^4$ unique SNPs (not found by the other methods) for each method.


```{r venndiagramsiggenes, echo=FALSE,fig.align="center", fig.cap="A Venn diagram of the overlaps in SNP for RFlocalfdr, Boruta, AIR, RFE and PIMP. AIR and PIMP are very much the outliers, with more than $10^4$ unique SNPs (not found by the other methods) for each method.", out.width = '50%'}
knitr::include_graphics("./figs/ven_diagram_sig_genes.png")

```


# References

    
```{r ,echo=FALSE,eval=TRUE}
knitr::knit_exit()

```

job_id=$(sbatch --parsable test.sh)
echo $job_id
17211434

sacct --starttime 2023-04-24 --format="jobid,Elapsed,TotalCPU,MaxRSS,AveVMSize,MaxVMSize,ReqMem"
dd:hh:mm:ss

166467464
169469556
###################################################
script20  Boruta  
slurmstepd: error: *** JOB 22009163 ON c001 CANCELLED AT 2023-05-10T20:19:52 DUE TO TIME LIMIT ***
petrichor-i1: sbatch script20.q
Submitted batch job 21956889
etrichor-i1: sbatch script20.q
Submitted batch job 22009163
petr
etrichor-i1: sacct -j 22009163  --format="jobid,ntasks,Elapsed,TotalCPU,MaxRSS,AveVMSize,MaxVMSize,ReqMem"
       JobID   NTasks    Elapsed   TotalCPU     MaxRSS  AveVMSize  MaxVMSize     ReqMem 
------------ -------- ---------- ---------- ---------- ---------- ---------- ---------- 
22009163                05:00:21  00:00.080                                       200Gn 
22009163.ba+        1   05:00:29  00:00.079 158957016K 160208656K 160208656K      200Gn 
22009163.ex+        1   05:00:21  00:00.001       511K    152208K    152208K      200Gn 
pet


####################################################
script21  AIR         125322.943    256.062    2976.523 
                      1:10:48:42                  00:49:36

 user    system   elapsed 
86781.238    46.470  3000.075 


petrichor-i1: sacct -j 21185985  --format="jobid,ntasks,Elapsed,TotalCPU,MaxRSS,AveVMSize,MaxVMSize,ReqMem"
       JobID   NTasks    Elapsed   TotalCPU     MaxRSS  AveVMSize  MaxVMSize     ReqMem 
------------ -------- ---------- ---------- ---------- ---------- ---------- ---------- 
21185985                00:50:47 1-00:07:53                                       200Gn 
21185985.ba+        1   00:50:47 1-00:07:53  81500426K  82435008K  82435008K      200Gn 
21185985.ex+        1   00:50:49  00:00.002      1037K    152208K    152208K      200Gn 
##########
####################################################


script22  RFlocalfdr   

66619.705   163.805  1966.151 
30.267   0.552  30.821 
27.760   0.436  28.197 

66619.705+30.267+27.760 
 1966.151 +30.821 +  28.197 

etrichor-i1: sacct -j 21779830  --format="jobid,ntasks,Elapsed,TotalCPU,MaxRSS,AveVMSize,MaxVMSize,ReqMem"
       JobID   NTasks    Elapsed   TotalCPU     MaxRSS  AveVMSize  MaxVMSize     ReqMem 
------------ -------- ---------- ---------- ---------- ---------- ---------- ---------- 
21779830                00:57:54   18:36:06                                       200Gn 
21779830.ex+        1   00:57:55   00:00:00       179K    152208K    152208K      200Gn 
21779830.0          1   00:57:51   18:36:06  80748741K  83253836K  83253836K      200Gn 
pet

####################################################
script23  PIMP        out of memory again Submitted batch job 21238051 21257516


####################################################
script24  RFE          running ubmitted batch job 21241826     21272164      time out after 24 hours
                                                               
 sacct -j 21272164  --format="jobid,ntasks,Elapsed,TotalCPU,MaxRSS,AveVMSize,MaxVMSize,ReqMem"
 JobID   NTasks         Elapsed   TotalCPU     MaxRSS  AveVMSize  MaxVMSize     ReqMem 
21272164              1-00:00:05  00:00.072                                       100Gn 
21272164.ba+        1 1-00:00:06  00:00.070  79405500K  80928772K  80928772K      100Gn 
21272164.ex+        1 1-00:00:05  00:00.001       816K    152208K    152208K      100Gn 

21381999 asked for 4 days   --- did not finish CANCELLED AT 2023-05-02T19:53:14 DUE TO TIME LIMIT 
96 hours

####################################################
I think this was script24  RFE
"slurm-21201677.out" 1L, 99B                                                                     1,1           All
slurmstepd: error: *** JOB 21201677 ON c349 CANCELLED AT 2023-04-24T21:49:19 DUE TO TIME LIMIT ***
etrichor-i1: sacct -j 21201677  --format="jobid,ntasks,Elapsed,TotalCPU,MaxRSS,AveVMSize,MaxVMSize,ReqMem"
       JobID   NTasks    Elapsed   TotalCPU     MaxRSS  AveVMSize  MaxVMSize     ReqMem 
------------ -------- ---------- ---------- ---------- ---------- ---------- ---------- 
21201677                05:00:17  00:00.073                                       200Gn 
21201677.ba+        1   05:00:21  00:00.071  78698997K  80928772K  80928772K      200Gn 
21201677.ex+        1   05:00:17  00:00.002      1039K    152208K    152208K      200Gn 


I think this was script23  PIMP
slurm-21206043.out
out-of-memory handler
petrichor-i1: sacct -j 21206043  --format="jobid,ntasks,Elapsed,TotalCPU,MaxRSS,AveVMSize,MaxVMSize,ReqMem"
       JobID   NTasks    Elapsed   TotalCPU     MaxRSS  AveVMSize  MaxVMSize     ReqMem 
------------ -------- ---------- ---------- ---------- ---------- ---------- ---------- 
21206043                05:00:31  00:00.075                                       600Gn 
21206043.ba+        1   05:01:10  00:00.074 633289727K 124075959+ 124075959+      600Gn 
21206043.ex+        1   05:00:31  00:00.001       572K    152208K    152208K      600Gn 


 sacct -j 21185758  --format="jobid,ntasks,Elapsed,TotalCPU,MaxRSS,AveVMSize,MaxVMSize,ReqMem"
 sacct -j 21185985  --format="jobid,ntasks,Elapsed,TotalCPU,MaxRSS,AveVMSize,MaxVMSize,ReqMem"
21185985

etrichor-i1: sacct --starttime 2023-04-24 --format="jobid,Elapsed,TotalCPU,MaxRSS,AveVMSize,MaxVMSize,ReqMem"
       JobID    Elapsed   TotalCPU     MaxRSS  AveVMSize  MaxVMSize     ReqMem 
------------ ---------- ---------- ---------- ---------- ---------- ---------- 
21185735       00:00:46  00:44.206                                       200Gn 
21185735.ba+   00:00:46  00:44.204   7282773K   7420288K   7420288K      200Gn 
21185735.ex+   00:00:46  00:00.002      1046K    152208K    152208K      200Gn 
21185747       00:00:01  00:00.073                                       200Gn 
21185747.ba+   00:00:01  00:00.073      1298K    152556K    152556K      200Gn 
21185747.ex+   00:00:01   00:00:00       540K    152208K    152208K      200Gn 
21185758       03:15:59   09:01:59                                       200Gn 
21185758.ba+   03:15:59   09:01:59 166467464K 169469556K 169469556K      200Gn 
21185758.ex+   03:15:59  00:00.002      1046K    152208K    152208K      200Gn 
21185806       00:02:04  02:04.681                                       200Gn 
21185806.ba+   00:02:04  02:04.678  76967681K  77585372K  77585372K      200Gn 
21185806.ex+   00:02:04  00:00.002       864K    152208K    152208K      200Gn 
21185985       00:50:47 1-00:07:53                                       200Gn 
21185985.ba+   00:50:47 1-00:07:53  81500426K  82435008K  82435008K      200Gn 
21185985.ex+   00:50:49  00:00.002      1037K    152208K    152208K      200Gn 
21190451       00:35:20   19:22:37                                       200Gn 
21190451.ba+   00:35:20   19:22:37  80922258K  82435008K  82435008K      200Gn 
21190451.ex+   00:35:20  00:00.001       739K    152208K    152208K      200Gn 
21196957       00:33:25   18:11:35                                       200Gn 
21196957.ba+   00:33:25   18:11:35  80731538K  82435008K  82435008K      200Gn 
21196957.ex+   00:33:25  00:00.001       711K    152208K    152208K      200Gn 
21197335       00:35:20   17:36:38                                       200Gn 
21197335.ba+   00:35:20   17:36:38 206690301K 106100719+ 106100719+      200Gn 
21197335.ex+   00:35:20  00:00.001       958K    152208K    152208K      200Gn 
21200418       00:01:46  01:44.233                                       200Gn 
21200418.ba+   00:01:46  01:44.232  22483114K  22685672K  22685672K      200Gn 
21200418.ex+   00:01:46  00:00.001       995K    152208K    152208K      200Gn 
21201677       05:00:17  00:00.073                                       200Gn 
21201677.ba+   05:00:21  00:00.071  78698997K  80928772K  80928772K      200Gn 
21201677.ex+   05:00:17  00:00.002      1039K    152208K    152208K      200Gn 
21202167       00:35:05   17:10:57                                       400Gn 
21202167.ba+   00:35:05   17:10:57 416721903K 107580308+ 107580308+      400Gn 
21202167.ex+   00:35:05  00:00.001      1162K    152208K    152208K      400Gn 
21206043       05:00:31  00:00.075                                       600Gn 
21206043.ba+   05:01:10  00:00.074 633289727K 124075959+ 124075959+      600Gn 
21206043.ex+   05:00:31  00:00.001       572K    152208K    152208K      600Gn 
21238051       05:00:14 2-00:31:03                                   1019390Mn 
21238051.ba+   05:00:57 2-00:31:03 103902005+ 158715088+ 158715088+  1019390Mn 
21238051.ex+   05:00:16   00:00:00       542K    152208K    152208K  1019390Mn 
21241826       10:00:30  00:00.075                                       200Gn 
21241826.ba+   10:00:31  00:00.073  78934632K  80928772K  80928772K      200Gn 
21241826.ex+   10:00:30  00:00.001       578K    152208K    152208K      200Gn 
21257516       00:00:00   00:00:00                                      2000Gn 
21272164       00:31:03   00:00:00                                       100Gn 
21272164.ba+   00:31:03   00:00:00                                       100Gn 
21272164.ex+   00:31:03   00:00:00                                       100Gn 


###############################################################################################

/scratch1/dun280/chr22_phase3_dpas03/script30

script20  Boruta  
etrichor-i1: sbatch script20.q
Submitted batch job 21959506
petrichor-i1: sbatch script20.q
Submitted batch job 22006019
petrichor-i1: 


slurmstepd: error: *** JOB 22006019 ON c026 CANCELLED AT 2023-05-10T20:07:38 DUE TO TIME LIMIT ***
etrichor-i1: sacct -j 22006019  --format="jobid,ntasks,Elapsed,TotalCPU,MaxRSS,AveVMSize,MaxVMSize,ReqMem"
       JobID   NTasks    Elapsed   TotalCPU     MaxRSS  AveVMSize  MaxVMSize     ReqMem 
------------ -------- ---------- ---------- ---------- ---------- ---------- ---------- 
22006019                05:00:26  00:00.066                                       200Gn 
22006019.ba+        1   05:00:34  00:00.065 166791307K 169454840K 169454840K      200Gn 
22006019.ex+        1   05:00:26  00:00.001       570K    152208K    152208K      200Gn 
pe


     user    system   elapsed 
32783.848  2427.226 11763.086
petrichor-i1: sacct -j 21937889  --format="jobid,ntasks,Elapsed,TotalCPU,MaxRSS,AveVMSize,MaxVMSize,ReqMem"
       JobID   NTasks    Elapsed   TotalCPU     MaxRSS  AveVMSize  MaxVMSize     ReqMem 
------------ -------- ---------- ---------- ---------- ---------- ---------- ---------- 
21937889                03:16:51   09:47:38                                       200Gn 
21937889.ba+        1   03:16:51   09:47:38 153749758K 156635004K 156635004K      200Gn 
21937889.ex+        1   03:16:51  00:00.001       569K    152208K    152208K      200Gn 


script21  AIR 
   user   system  elapsed
9940.345   85.390  359.682
petrichor-i1: sacct -j 21937892  --format="jobid,ntasks,Elapsed,TotalCPU,MaxRSS,AveVMSize,MaxVMSize,ReqMem"
       JobID   NTasks    Elapsed   TotalCPU     MaxRSS  AveVMSize  MaxVMSize     ReqMem 
------------ -------- ---------- ---------- ---------- ---------- ---------- ---------- 
21937892                00:06:46   02:47:50                                       200Gn 
21937892.ba+        1   00:06:46   02:47:50  77358209K  79836120K  79836120K      200Gn 
21937892.ex+        1   00:06:46  00:00.001       544K    152208K    152208K      200Gn 


script22  RFlocalfdr 
petrichor-i1: sbatch script22.q
Submitted batch job 21948338
sacct -j 21948338  --format="jobid,ntasks,Elapsed,TotalCPU,MaxRSS,AveVMSize,MaxVMSize,ReqMem"
      JobID   NTasks    Elapsed   TotalCPU     MaxRSS  AveVMSize  MaxVMSize     ReqMem 
------------ -------- ---------- ---------- ---------- ---------- ---------- ---------- 
21948338                00:05:15   01:49:34                                       200Gn 
21948338.ba+        1   00:05:15   01:49:34  77306709K  79836144K  79836144K      200Gn 
21948338.ex+        1   00:05:15  00:00.001       842K    152208K    152208K      200Gn 
petrichor-i1: 
   user   system  elapsed 
6452.098   72.372  261.622 
  3.199   0.056   3.258 
  1.100   0.096   1.365 
  0.043   0.000   0.066 

petrichor-i1: sbatch script22.q
Submitted batch job 22081543


script23  PIMP   
petrichor-i1: vi script23.txt 
petrichor-i1: rm script23.txt 
rm: remove regular file 'script23.txt'? y
petrichor-i1: sbatch script23.q
Submitted batch job 21948345
outof memory
etrichor-i1: sacct -j 21948345  --format="jobid,ntasks,Elapsed,TotalCPU,MaxRSS,AveVMSize,MaxVMSize,ReqMem"
       JobID   NTasks    Elapsed   TotalCPU     MaxRSS  AveVMSize  MaxVMSize     ReqMem 
------------ -------- ---------- ---------- ---------- ---------- ---------- ---------- 
21948345                00:07:13   01:58:53                                       200Gn 
21948345.ba+        1   00:07:13   01:58:53 208149440K 106114068+ 106114068+      200Gn 
21948345.ex+        1   00:07:13  00:00.001       734K    152208K    152208K      200Gn 
petrichor-i1: 


script24  RFE    
petrichor-i1: vi script24.txt 
petrichor-i1: rm script24.txt 
rm: remove regular file 'script24.txt'? y
petrichor-i1: sbatch script24.q
Submitted batch job 21948351
petrichor-i1: sacct -j 21948351  --format="jobid,ntasks,Elapsed,TotalCPU,MaxRSS,AveVMSize,MaxVMSize,ReqMem"


#############################################################################################################
/scratch1/dun280/chr22_phase3_dpas03/script21


\begin{table}[tbhp]
  \centering
  \begin{tabular}{||p{2cm}|l|D{.}{.}{3}|D{.}{.}{3}|D{.}{.}{3}|c||}
    \hline
    Method     &  SNPSs returned&\multicolumn{1}{c|}{elapsed}    &\multicolumn{1}{c|}{user} &  \multicolumn{1}{c|}{MaxRSS (GB)}   &notes           \\
    \hline
    RFlocalfdr & 6335          &   2025.17   &  66677.73     &  79.40       &               \\
    Boruta     & -             &  -          &  -            &              &  out of time  \\
    RFE        & -             &  345600.00  &   -           &   -          &               \\
    AIR        & 191489        &  2976.52    & 125322.94     &  77.72       &               \\
    PIMP       &  -            &     -       &    -          &       -      & out of memory    \\
    \hline
  \end{tabular}
     \caption{
    A more systematic, but still somewhat arbitrary, attempt to measure the run-times. The files for replicating this
   trial are included in the **RFlocalfdr** package.  RFE dd not complete in 96 hours. PIMP failed many times because of
   memory limits. We progressively increased the memory to 2000GB. However, due to the size of the requested
   resources, this failed to run for over a week. }
    \label{systematic_thousand_genome_results_100000}
\end{table}


#script40
petrichor-i1: sbatch script20.q
Submitted batch job 22055833
petrichor-i1: sbatch script21.q
Submitted batch job 22055834
petrichor-i1: sbatch script22.q
Submitted batch job 22055836
petrichor-i1: sbatch script23.q
Submitted batch job 22055837
petrichor-i1: sbatch script24.q
Submitted batch job 22055838

petrichor-i1: sbatch script22.q
Submitted batch job 22081428


#junk
\begin{table}[tbhp]
  \centering
  \begin{tabular}{||p{2cm}|l|D{.}{.}{3}|D{.}{.}{3}|D{.}{.}{3}|c||}
    \hline
    Method     &  SNPSs returned&\multicolumn{1}{c|}{elapsed}    &\multicolumn{1}{c|}{user} &  \multicolumn{1}{c|}{MaxRSS (GB)}   &notes           \\
    \hline
    RFlocalfdr & 6335          &   2025.17   &  66677.73     &  79.40       &               \\
    Boruta     & -             &  -          &  -            &              &  out of time  \\
    RFE        & -             &  -           &   -           &   -         &  out of time  \\
    AIR        & 191489        &  2976.52    & 125322.94     &  77.72      &               \\
    PIMP       &  -            &     -       &    -          &       -      & out of memory    \\
    \hline
  \end{tabular}
     \caption{
    A more systematic, but still somewhat arbitrary, attempt to measure the run-times. The files for replicating this
   trial are included in the **RFlocalfdr** package.  RFE dd not complete in 96 hours. PIMP failed many times because of
   memory limits. We progressively increased the memory to 2000GB. However, due to the size of the requested
   resources, this failed to run for over a week. }
    \label{systematic_thousand_genome_results_100000}
\end{table}


\begin{table}[tbhp]
  \centering
  \begin{tabular}{||l|p{2cm}|l|D{.}{.}{3}|D{.}{.}{3}|D{.}{.}{3}|c||}
    \hline
    Method     &   ntree  & SNPSs returned&\multicolumn{1}{c|}{elapsed}    &\multicolumn{1}{c|}{user} &  \multicolumn{1}{c|}{MaxRSS (GB)}   &notes           \\
    \hline
    RFlocalfdr & 10000& 753          &    263.311   &   6456.44     &  79.40       &               \\
    Boruta     & 10000& -             &  -          &  -            &   161.60     &  out of time  \\
    RFE        &  10000& -             &  -           &   -          &   -         &  out of time  \\
    AIR        &  10000&155154        &  2976.52    & 9940.345       &  76.13      &               \\
    PIMP       &  10000& -            &     -       &    -          &       -      & out of memory    \\
    \hline
  \end{tabular}
     \caption{
    A more systematic, but still somewhat arbitrary, attempt to measure the run-times. The files for replicating this
   trial are included in the **RFlocalfdr** package.  RFE dd not complete in 96 hours. PIMP failed many times because of
   memory limits. We progressively increased the memory to 2000GB. However, due to the size of the requested
   resources, this failed to run for over a week. }
    \label{systematic_thousand_genome_results_10000}
\end{table}

Figure \ref{systematic_thousand_genome_results_10000}

- 40 cores, 200GB of RAM and 5 hours
- num.trees set to 10000 


\begin{table}[tbhp]
  \centering
  \begin{tabular}{||l|p{2cm}|l|D{.}{.}{3}|D{.}{.}{3}|D{.}{.}{3}|c||}
    \hline
    Method     &  ntree & SNPSs returned&\multicolumn{1}{c|}{elapsed}    &\multicolumn{1}{c|}{user} &  \multicolumn{1}{c|}{MaxRSS (GB)}   &notes           \\
    \hline
    RFlocalfdr &  1000 & 580         &  92.73     & 739.057     &  76.11        &         \\
    Boruta     & 1000 &  12081       & 12442.77   & 67078.45    &  66.21   &   \\
    RFE        & 1000 &              &            &             &          &   out of time\\
    AIR        & 1000 & 88225        &  107.88    &1114.116     &  76.11   &    \\
    PIMP       &  1000 &  -          &    -         &  -           & -         & out of memory  \\
    \hline
  \end{tabular}
     \caption{
    A more systematic, but still somewhat arbitrary, attempt to measure the run-times. The files for replicating this
   trial are included in the **RFlocalfdr** package.  RFE dd not complete in 96 hours. PIMP failed many times because of
   memory limits. We progressively increased the memory to 2000GB. However, due to the size of the requested
   resources, this failed to run for over a week. }
    \label{systematic_thousand_genome_results_1000}
\end{table}

Figure \ref{systematic_thousand_genome_results_1000}

- 40 cores, 200GB of RAM and 5 hours
- num.trees set to 10000 


\begin{table}[tbhp]
  \centering
  \begin{tabular}{||p{2cm}|l|l|l|l|l|l|l||}
    \hline
    Method     & SNPs returned & run time  \\
    \hline
    RFlocalfdr & 6,335          & 1       \\
    Boruta     & 6,773          & 100        \\
    RFE        & 12             & 5       \\
    AIR        & 61,092         & 2      \\
    PIMP       & 787502         & 100      \\
    \hline
  \end{tabular}
  \caption{The number of SNPs returned by each methods and the run time. Run times are expressed in multiples of the run time
    of a single ranger fit for the given hardware configuration, so we are really looking at the number of refits each
    method requires. The processing time outside of the RF fit is negligible, by comparison, in all cases.
	The RFE time is greater than 5 (see table \ref{systematic_thousand_genome_results}, but we were unable to detemine by how much. 
  \label{notional_thousand_genome_results}}
\end{table}
````
